# LIGR-Nue for the global mapping of the bacterial sRNA targetome *in vivo*

**DOI:** 10.64898/2026.11.09.765264

**Authors:** Mukulika Mandal, Christopher Adelmann, Jens Georg, Sreenivasulu Kurukuti, Björn Voß, Wolfgang R Hess, Jogadhenu S S Prakash

## Abstract

Interactions between regulatory small RNAs (sRNAs) and their target transcripts are central for post-transcriptional regulation in bacteria. Based on RNA-seq analysis, we determined the relative abundance of several sRNAs in the photosynthetic cyanobacterium *Synechocystis* sp. PCC 6803 under photoautotrophic growth conditions and after transfer to darkness. We observed an increase in the abundance of the sRNAs DsiR1 to 58.6% and of CsiR1 to 19% of the sRNA transcriptome after transfer to darkness. We then developed a duplex-directed, transcriptome-wide mapping strategy of RNA–RNA interactions *in vivo*, called LIGR-Nue, which involves aminomethyltrioxsalen-mediated UV cross-linking, proximity ligation, crosslink reversal, reverse transcription and sequencing. The data were analysed using the RNANUE pipeline for the detection of chimeric reads and mapping, and benchmarked against known targetome data for the sRNAs PsrR1 and NsiR4. Using this methodology, we identified RNA–RNA interactions, such as PsrR1 as a regulator of carbon assimilation, and DsiR1 as a global silencing factor of photosynthesis-associated mRNAs in the dark. This data enables the comprehensive analysis of sRNA functions in post-transcriptional gene regulation. LIGR-Nue is broadly applicable to map RNA–RNA interactions in bacteria.

**GRAPHICAL ABSTRACT:** 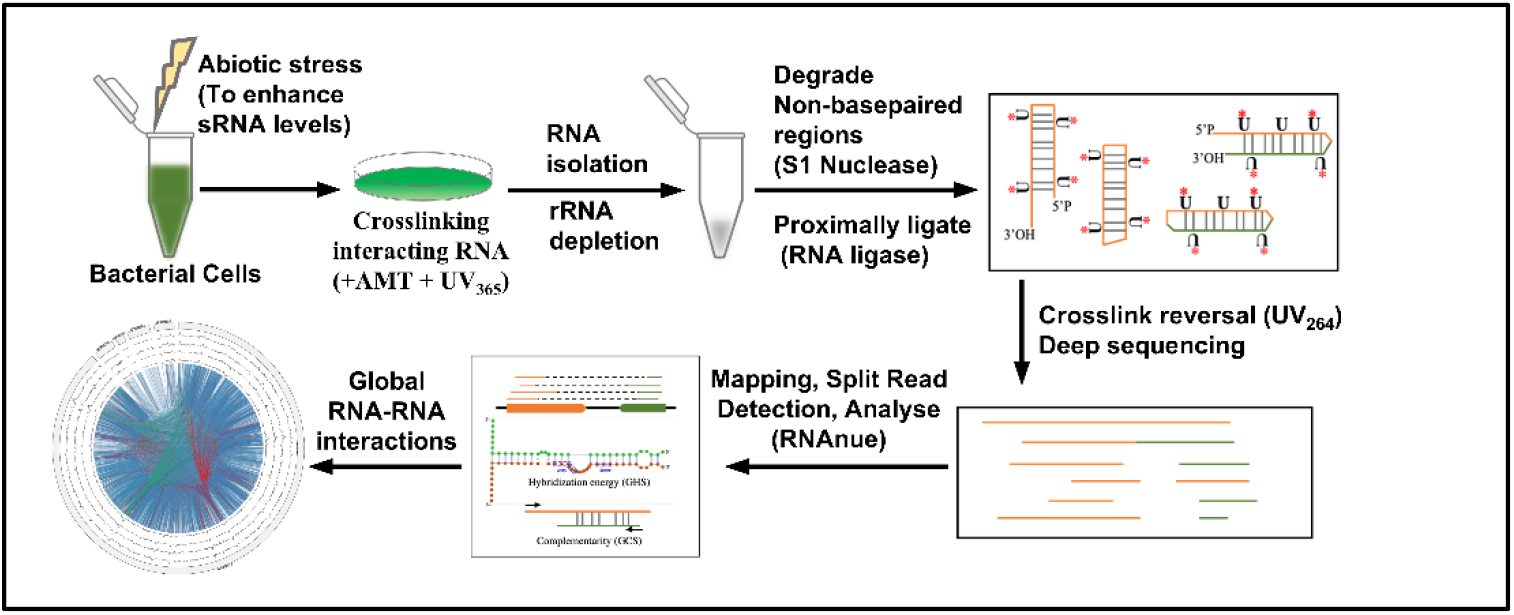

## INTRODUCTION

Bacteria have evolved complex regulatory networks to adapt to fluctuating environmental conditions and diverse abiotic and biotic stresses. An important element in these networks are non-coding RNAs (ncRNAs). Their roles are considerably less well characterized than those of regulatory proteins, primarily due to their lower level of conservation and the various technical challenges involved. While ncRNA functions are comparatively better studied in a few model bacteria, particularly in pathogens such as *Salmonella* [1], or *Staphylococcus aureus* [2], they remain largely understudied in most other bacterial taxa. Cyanobacteria rely on light as the primary energy source, occupy highly diverse ecological niches and show an exceptional morphological, metabolic, and regulatory versatility. These metabolic and adaptive capabilities have also positioned cyanobacteria as attractive platforms for biotechnology, as they can efficiently convert solar energy and carbon dioxide into valuable products such as biofuels, pigments and diverse secondary metabolites with pharmaceutical potential [3]. Fully exploiting this potential requires a detailed understanding of the regulatory networks governing cyanobacterial physiology and adaptability. This includes post-transcriptional control mediated by ncRNAs.

One class of ncRNAs are cis-encoded antisense RNAs (asRNAs), which are transcribed from the reverse strand of the gene they target, i.e. in cis [4]. Prominent examples of such asRNAs in the cyanobacterium *Synechocystis* sp. PCC 6803 (hereafter *Synechocystis*) include IsrR, which protects photosystem I (PSI) from photo-induced damage by regulating *isiA* expression [5]; PsbA2R and PsbA3R, which modulate *psbA* expression to optimize D1 protein synthesis [6]; RblR, which regulates the expression of the Rubisco large subunit (*rbcL*) and influences overall photosynthetic activity [7]; CoaR, involved in the regulation of coenzyme A biosynthesis and associated with butanol tolerance [8]; and PilR, which controls pilus gene expression and thereby affects cell motility [9].

However, the most important category of ncRNAs are small regulatory RNAs (sRNAs), which enable rapid, fine-tuned responses to environmental and metabolic cues [10]. In *Synechocystis*, dRNA-Seq analysis identified 371 non-coding transcriptional start sites (nTSS) for sRNA transcription, accounting for approximately 5.5% of all mapped TSS [11, 12]. Subsequently, it was found that many sRNAs are under tight regulatory control and display pronounced condition-dependent expression. Several sRNAs are undetectable under standard laboratory conditions, but are strongly and specifically induced in response to environmental stresses [11]. Among them, 33 abundant sRNAs were found to accumulate under 10 different abiotic stress conditions [12]. It is well documented that sRNAs frequently regulate gene expression by base-pairing with multiple, different target mRNAs under specific environmental or physiological conditions [13]. In *Synechocystis*, only a limited number of sRNAs have been functionally characterized so far. Several sRNAs in *Synechocystis* display strong condition-dependent expression patterns, suggesting important roles in environmental acclimation. For example, DsiR1 (also called PmgR1 and Ncr0700) [14] accumulates during darkness, stationary phase, and heat stress, while NsiR4, PsiR1, and IsaR1 are specifically induced under nitrogen, phosphate, and iron limitation, respectively. PsrR1 is strongly induced under high-light conditions. Furthermore, DsiR1, CsiR1, Ncr1220, and SyR52 exhibit expression patterns linked to inorganic carbon availability, a key factor for photosynthetic cyanobacteria [12,15–17]. The regulatory targets of some of these stress-responsive sRNAs have been identified in *Synechocystis*. The high-light inducible sRNA PsrR1 post-transcriptionally regulates genes encoding PSI and phycobilisome proteins in coordination with the response regulator RpaB [15,18,19], while the 3’end-derived sRNA ApcZ targets the *ocp* mRNA encoding the orange carotenoid protein to avoid physiological conflict between light harvesting and photoprotection [20].

Several sRNAs are induced under specific nutrient limitations. The nitrogen starvation induced sRNA NsiR4 controls expression of the glutamine synthetase inactivating factor A (GifA) [21], and of the regulatory protein PirA [22]. The most complex regulon was found to be controlled by the iron starvation-induced sRNA IsaR1 [17, 23]. While these examples illustrate that *Synechocystis* is the best-studied model for post-transcriptional regulation among cyanobacteria, many sRNAs that accumulate under abiotic stress conditions still lack identified RNA targets and defined regulatory roles. Therefore, establishing a global RNA– RNA interaction (RRI) network is essential to link sRNAs with their regulatory targets and to better understand post-transcriptional regulation mediated by sRNAs at a systems level.

Several methods have been developed to capture RRIs globally. Early approaches such as PARS [24], FragSeq [25], and PARTE [26] analyse RNA secondary structures *in vitro* using chemical or enzymatic probing; however, these methods rely on RNA extraction and refolding and therefore do not reflect native cellular conditions shaped by RNA-binding proteins (RBPs). Protein-centric techniques like CLIP identify RBP–RNA binding sites but cannot directly distinguish intra-from intermolecular RRIs [27]. To address this, methods such as CLASH [28], and later hiCLIP [29] and RIPPLiT [30], introduced proximity ligation of RNAs within RBP complexes, enabling detection of interacting RNAs as chimeric reads, although these approaches remain largely protein-centric [31].

To capture RRIs more broadly and under near-physiological conditions, chemical cross-linking–based methods were developed. One such method, LIGR-seq, uses psoralen-mediated cross-linking of RNA duplexes *in vivo* followed by proximity ligation and high-throughput sequencing and was applied to identify transcriptome-wide interacting RNA pairs in humans [32]. More recently, a bioinformatic pipeline, RNANUE, has been developed for inferring RRIs from raw Direct Duplex Data (DDD) [33]. RNANUE integrates RNA hybridization potential, thermodynamic stability, and rigorous statistical evaluation, thereby improving the sensitivity and specificity of detecting RRIs. Importantly, it also enables the detection of interactions involving previously unannotated transcripts, making it particularly useful for studying RNA regulatory networks in non-model organisms [33].

In this study, we developed LIGR-Nue, an experimental approach to identify RRIs globally in the cyanobacterium *Synechocystis in vivo.* For this, we adapted the LIGR-seq workflow originally developed for human cell lines, with specific modifications, followed by analysis using the RNANUE bioinformatics pipeline. To our knowledge, this is the first application of a duplex-directed, transcriptome-wide RRI mapping strategy in cyanobacteria, providing a global snapshot of the RNA interactome in this photosynthetic model organism. This systems-level approach enabled the identification of numerous previously uncharacterized RRIs. A major focus of this study is the methodological development and benchmarking of this strategy in *Synechocystis,* thereby establishing a framework for future investigations of RNA-mediated regulation in cyanobacteria, photosynthetic microbes, and other bacteria.

## MATERIAL AND METHODS

### Culture conditions

A glucose-tolerant strain of *Synechocystis* [34], originally obtained from Dr J. G. K. Williams (Dupont de Nemours) was photoautotrophically (PA) grown at 34 °C in BG-11 medium [35], buffered with 20 mM HEPES/NaOH (pH 7.5), under continuous illumination from incandescent lamps at 70 µmol photons m^−2^ s^−1^, as described previously [36]. For dark and high light treatment, cultures grown photoautotrophically to mid exponential phase (∼0.5 at OD_730nm_) were shifted to dark for 1 h (dark), or illuminated at 400 µmol photons m^−2^ s^−1^ for 30 minutes (HL), respectively.

### RNA extraction and cDNA library preparation for whole transcriptome analysis (WTA)

S*ynechocystis* cells grown under PA and dark in duplicates were fixed instantaneously by the addition of 50 mL ice-cold phenol/ethanol (1 : 20, w/v) to 50 mL of the cell suspension and then total RNA was extracted as described previously [37]. The RNA samples were quantitated on Qubit® fluorometer and appropriate dilutions loaded on a High sensitivity RNA screen tape to determine the RNA integrity number (RIN). Briefly, 1 μg of QC-verified RNA samples was depleted of rRNA using the NEBNext® rRNA depletion kit (Bacteria, Cat. no. E7850), according to the manufacturer’s protocol. This included a DNase I digestion step. The rRNA-depleted RNA was purified using RNA purification beads as instructed. The NEBNext® Ultra II Directional RNA Library prep kit for Illumina (Cat. no. E7760L), was used to construct double-stranded cDNA libraries from the rRNA-depleted RNA, using RNA fragmentation, first strand cDNA synthesis with random primers, second strand cDNA synthesis, end repair of double stranded cDNA, adapter ligation, removal of excess adapter using sample purification beads, PCR enrichment of adapter ligated DNA and clean-up of PCR products as instructed by the manufacturer. The cleaned libraries were quantitated on Qubit® fluorometer and appropriate dilutions loaded on a High sensitivity D1000 screen tape to determine the size range of the fragments and the average library size. Sequencing was performed on an Illumina NovaSeqX Plus (Illumina Inc., San Diego, CA, USA) platform using paired end 150 bp sequencing. Data was demultiplexed using respective index sequences. Library preparation and sequencing was performed by Chromosome labs Pvt. Ltd (Mumbai, India).

### Bioinformatic pipeline for WTA

Raw reads (fastq) were initially assessed for quality using FastQC (https://www.bioinformatics.babraham.ac.uk/projects/fastqc/) to evaluate per-base sequence quality, GC content, and adapter contamination. Adapter sequences and low-quality bases were trimmed using Trimmomatic [38] removing bases with quality scores below 22 from both ends, discarding reads with average quality below 22, and retaining reads with a minimum length of 30 bp. The filtered high-quality reads were aligned to the *Synechocystis* genome (NCBI refseq assembly: GCF_000009725.1, ASM972v1) using Bowtie2 in end-to- end alignment mode [39]. Alignment output was generated in sequence alignment/map (SAM) format and were converted to compressed binary alignment/map format, sorted, and indexed using SAMtools [40], and only high-confidence alignments with mapping quality scores ≥30 (MAPQ ≥30) were retained for downstream analyses. Alignment quality and read distribution across genomic regions were visually inspected using IGV. Gene-level read counts were generated using BEDTools [41] by intersecting filtered BAM files with annotated gene coordinates. The reference genome annotation file (GFF format) obtained from the same NCBI assembly (GCF_000009725.1, ASM972v1) was manually curated to include sRNA features that were absent from the original annotation. Transcript abundance was normalized as transcripts per million (TPM) by calculating reads per kilobase (RPK) for each gene, summing all RPK values to obtain a scaling factor, and scaling individual RPK values to one million, enabling accurate comparison of gene expression levels across genes and samples.

### AMT crosslinking, RNA isolation and ribodepletion

*Synechocystis* cells grown under PA, dark, and HL were harvested and centrifuged at 2100 x g for 5 min at 4°C. The resulting pellet was resuspended in cold PBS followed by resuspension in 2 mL of ice-cold TS buffer. 4′-aminomethyltrioxsalen (AMT; Sigma, A4330) was added to a final concentration of 300 µg/mL. Cells were incubated on ice for 10 min with gentle mixing at 1-min intervals. The suspension was then transferred to an open 60-mm petri dish placed on ice and irradiated with 365 nm UV light for six cycles of 10 min each, maintaining a distance of 15 cm from the UV source and mixing the sample between irradiation cycles. Total RNA was extracted as previously described [37] and treated with TURBO DNase I (Thermo Fisher Scientific, AM1907) to remove genomic DNA contamination. Detailed experimental procedures are provided in the **Supplementary Material 1**.

### RNA processing and proximity ligation

Ribosomal RNA was depleted from 4 µg of total RNA using the NEBNext® rRNA Depletion Kit (Bacteria) (New England Biolabs, E7860). The resulting RNA was digested for 30 min at room temperature in a 40 µL reaction containing 1× S1 buffer and 2 µL S1 nuclease (diluted 1:100 in 1× S1 buffer; Thermo Fisher, EN0321). This was followed by NEBNext® RNA Sample Purification bead-based purification of the RNA (NEB, E7860). Ligation was performed using CircRNA ligase (Epicenter, CL4111K) in 20 µL reactions containing the entire S1-digested RNA. Samples were pre-incubated with 2 µL of 10× circRNA ligase buffer for 2 min at 85°C to denature the RNA and immediately transferred to ice. Subsequently, 1 µL of 10 mM ATP, 1 µL of 50 mM MnCl_₂_, and 1 µL of circRNA ligase (5 U/µL) were added, and the reactions were incubated in a thermocycler for 1 h at 60°C.

### Enrichment of crosslinked RNA and crosslink reversal

The reactions from the previous step were mixed with 0.5 µL of RNase R (Epicenter, RNR07250), 2.5 µL of RNase R buffer, and 2 µL of H₂O, and incubated for 10 min at 37°C. The reactions were then stopped by placing the tubes on ice. RNA was purified using bead-based extraction method, washed with 80% ethanol, and resuspended in 20 µL of TS buffer. Samples were transferred to open 2 mL microcentrifuge tubes, placed on ice, and irradiated at 254 nm for 10 min at a distance of 15 cm from the UV source, with 5-min interval between exposures (**Supplementary Material 1**).

### Library preparation and sequencing of chimeric reads

Total volume of RNA sample was used for library preparation. The NEBNext® Ultra II Directional RNA Library prep kit for Illumina (Cat. no. E7760L), was used to construct double-stranded cDNA libraries using first strand cDNA synthesis with random primers, second strand cDNA synthesis, end repair of double stranded cDNA, adapter ligation, removal of excess adapter using sample purification beads, PCR enrichment of adapter ligated DNA and clean-up of PCR products as instructed by the manufacturer. The cleaned libraries were quantitated on Qubit® fluorometer and appropriate dilutions loaded on a D1000 screen tape to determine the size range of the fragments and the average library size. Sequencing was performed on the Illumina NovaSeqX Plus (Illumina Inc., San Diego, CA, USA) platform using paired end 150 bp sequencing. Library preparation and sequencing was performed by Chromosome labs Pvt. Ltd (Mumbai, India).

### Bioinformatic pipeline for chimeric read detection and statistical analysis

LIGR-Seq fastq files were analysed using RNAnue v1.0.0-beta.1 (https://github.com/RNABioInfo/RNAnue/releases/tag/v1.0.0-beta.1) [33]. This includes a preprocessing step covering deduplication, read trimming (--wsize 1), paired-end read merging, and quality/length filtering (--minqual 10, --minlen 10). Alignment was performed using the segemehl algorithm (v. 0.3.4) [42] for chimeric read alignment (--minfragsco 10; GenBank: GCA_000009725.1, ASM972v1). Chimeric reads were filtered during detection using minimum complementarity (--sitelenratio 0.5; -- cmplmin 0.3), alignment length (-- mindetectlen 10), and hybridization energy (ViennaRNA v2.6.4). Reads were annotated by gff-annotation overlap (**Supplementary Material 3**) or assigned to supplementary features when no feature was present. Subsequently, chimeric reads were pairwise-clustered (-- clustmethod specific, --clustfrac 0.3) and count-aggregated by their respective scores. Furthermore, global complementarity and hybridization scores were calculated for each interaction cluster. Clusters with scores below five were discarded, and the remaining clusters were statistically evaluated against background feature counts. During postprocessing, interactions among samples within the same condition were aggregated into superclusters (--intfrac 0.5). Interacting regions were annotated according to genomic features (NCBI RefSeq assembly: GCA_000009725.1, ASM972v1) supplemented with transcription unit annotations (**Supplementary Material 3**). The Interaction result file contains metrics for each detected RNA–RNA interaction, including total_span_bp, effective_coverage_span_bp, support_per_total_bp, support_per_effective_bp, coverage_concentration, coverage_components, arm_balance, coverage_profile, no_splits, mean_inter_crosslinks, GCS, GHS and p-value which are described in **Supplementary Table 1**. The interaction network was constructed using the predicted targets of each sRNA, with node size scaled according to degree and edge width proportional to the IntaRNA interaction energy. Binding regions were visualized using SnapGene (Dotmatics, https://www.snapgene.com/). Interaction energies between the fragments detected by LIGR-Nue as chimeric reads were predicted using IntaRNA [43], and RNA secondary structures were predicted using RNAfold [44] from the ViennaRNA Package and visualized using VARNA [45]. Circos plots were generated using R, and interaction network was constructed and visualized using Cytoscape [46].

## RESULTS

### Abundance of small RNAs under darkness

In *Synechocystis*, sRNAs constitute less than 1% of the total cellular RNA, although some become strongly and specifically induced under particular abiotic stress conditions [47]. To increase the likelihood of detecting the sRNAs and their target RNA interactions *in vivo*, we focused on conditions that promote sRNA accumulation. Among the reported stress-responsive sRNAs, DsiR1 is known to accumulate under multiple stresses, particularly darkness. Therefore, we first assessed the relative abundance of sRNAs by whole-transcriptome analysis under optimal growth conditions (photoautotrophy, PA) and light-to-dark shift for 1 h (Dark) (**Figure 1**). The shift to darkness was chosen because cyanobacterial physiology in the dark is less understood compared to many other conditions. The total sRNA abundance increased markedly after 1 h in the dark, with Transcript Per Million (TPM) values rising from 17,518 ± 233 in PA to 75,800 ± 514, representing a 4.3-fold increase. Notably, the share of DsiR1 in the total sRNA reads increased dramatically, from 4.3% of under PA to 58.6%. Similarly, CsiR1 abundance increased from 1.35% under PA to 19% under dark (**Figure 1**). Other sRNAs with substantial TPM values under both conditions included SyR50, Tu818, DsiR3, SyR53, SyR7, SyR47, CoiR1, TU2529, SyR52, and SyR9. Based on the strong enrichment of several sRNAs, dark stress was selected for identifying sRNA–mRNA interactions, with PA serving as the control condition. In addition, since the mRNA targets of the sRNA PsrR1 under high light were previously characterized [15], high-light exposure for 30 min (HL) was included as an internal validation of the experimental approach.

**Figure 1:**
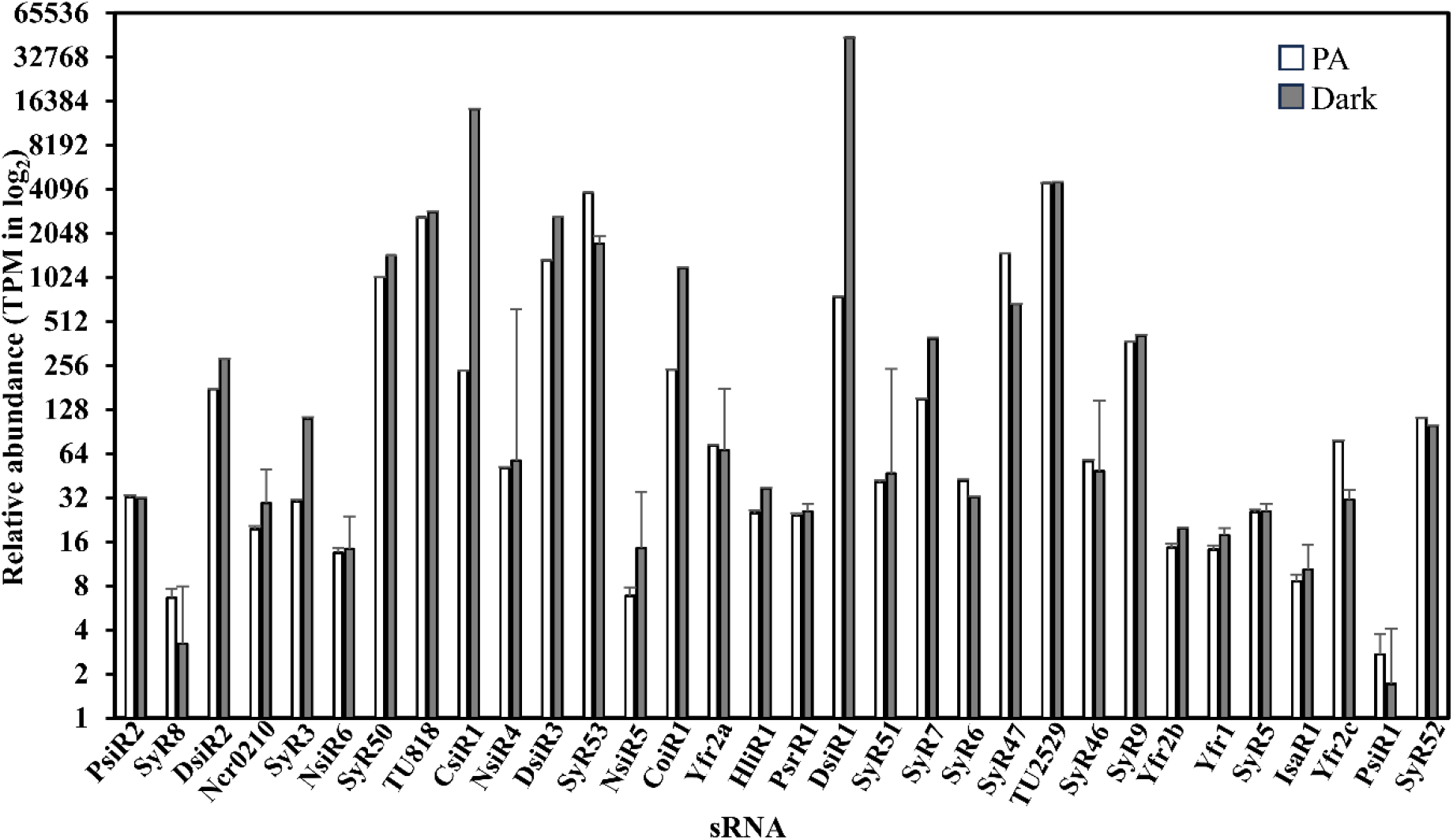
The relative abundance of 32 previously defined sRNAs (Kopf and Hess, 2015) under photoautotrophic growth and darkness. Photoautotrophically (PA) grown cells (OD_730nm_ = 0.5) were shifted to darkness for 1 h (Dark). RNA seq data of PA and Dark were generated in duplicates. sRNA abundances are represented as Transcripts per Million (TPM) on log_2_ scale. TPM of DsiR1 under PA is 754.7 ± 192 and 44,427.8 ± 195 under Dark. Similarly, CsiR1 has TPM values 236.5 ± 54 and 14,402 ± 566 under PA and Dark respectively. Note that NsiR6 and HliR1 have meanwhile been identified to encode the small proteins NblD and SliP4, respectively [48,49].

### Quality assessment of LIGR-Nue libraries and the pipeline

To systematically map sRNA–mRNA interactions, we optimized the duplex-directed RNA interaction mapping strategy based on LIGR-seq [32]. PA, HL and dark-treated samples were subjected to AMT crosslinking, proximity ligation, enrichment, and library preparation for Illumina sequencing (**Figure 2**). The detailed protocol is provided as **Supplementary Material 1**. To assess background ligation artifacts, control samples without AMT cross-linking were prepared and analysed in parallel. Tape Station profiles demonstrated consistent library quality across samples, with average fragment sizes ranging from ∼257– 486 bp. Marker peaks were clearly detected, and adapter-dimer contamination was not observed, indicating good library quality and uniformity across libraries (**Supplementary Figure S1**). All analysed ten libraries yielded high-quality paired-end reads (∼2–4.5 million paired-end reads per library) with a mean read length of 151 bp, >90% of bases at Q20, and >85% at Q30. The resulting high-throughput sequencing data were analysed using RNAnue v1.0.0-beta.1 (https://github.com/RNABioInfo/RNAnue/releases/tag/v1.0.0-beta.1) [33]. Raw reads generated from samples without crosslinking and ligation were also analysed. As expected, no chimeric RRIs were detected, confirming that the chimeras observed in LIGR-Nue libraries depended on the ligation step. Interestingly, a single split read observed in the ligation-negative control corresponded to the exons of the initiator tRNA, tRNA_i_^fmet^. The tRNA_i_^fmet^ primary transcript consists of two exons separated by a group I intron in which gene *slr0915* is located [50]. This chimera was not observed in AMT cross-linked and proximity-ligated samples, as it arises from exon fusion rather than RNA hybridization. This observation further supported the validity of the experimental strategy used to generate the sRNA–mRNA interactome in *Synechocystis* (**Figure 3**).

**Figure 2.**
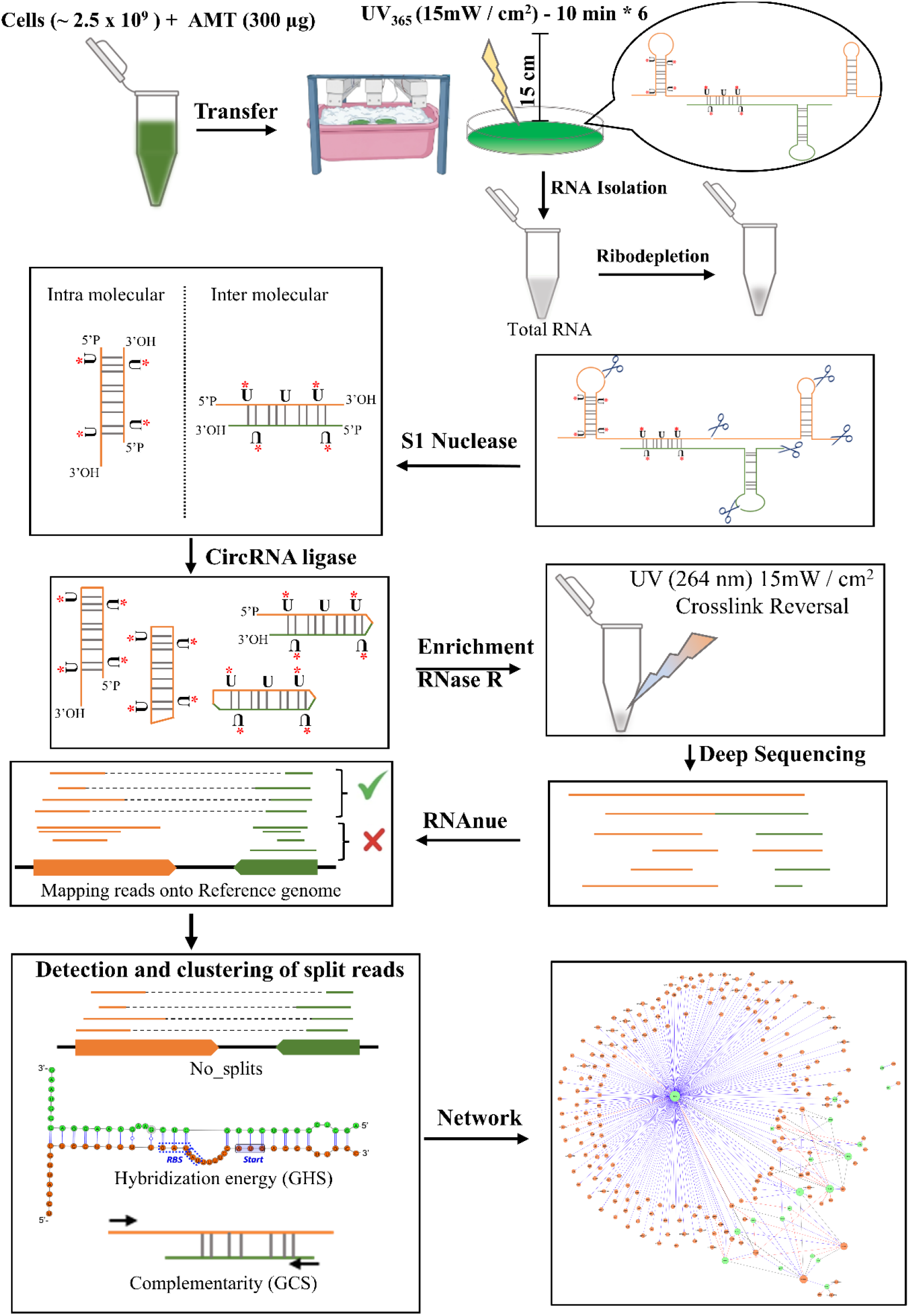
Graphical representation of the LIGR-Nue method.

**Figure 3.**
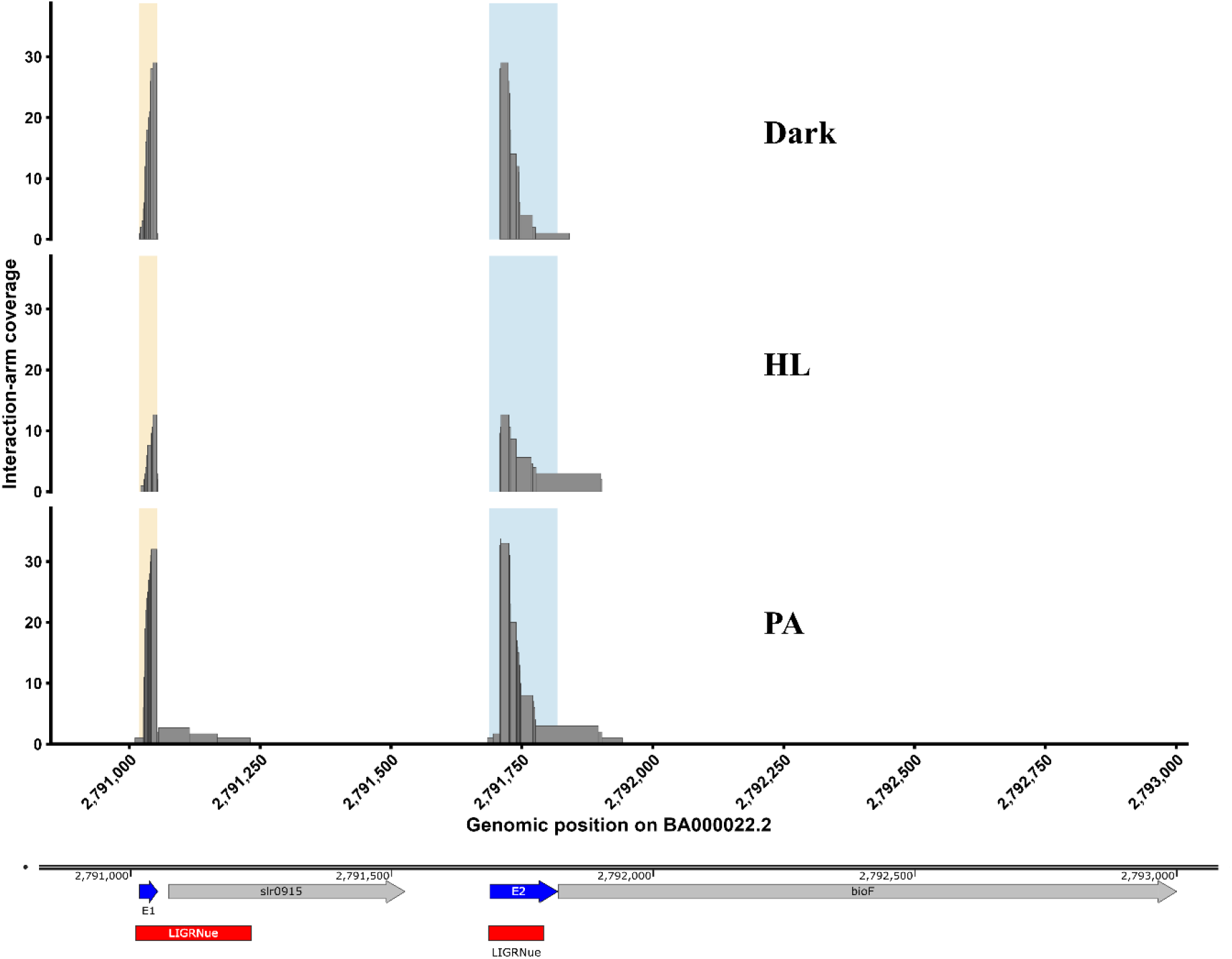
A spliced transcript illustrates a chimeric interaction between the two tRNA_i_^fmet^ exons. This tRNA originates from a single gene in *Synechocystis* and is interrupted by a group I intron just 5’ of the anticodon. The two exons, designated E1 and E2, were identified as a chimeric interaction in non-AMT-treated, non-ligated samples. Interaction-arm coverage across the E1–E2 genomic region is shown for cells grown under photoautotrophic conditions (PA), 1 h of dark (Dark), and 30 min of high light (HL). Shaded regions indicate the positions of E1 and E2, while the bars represent the corresponding interaction-arm coverage. The genomic organization of the locus, including *slr0915* and *bioF*, is shown below. The *slr0915* gene encodes an intron-encoded PD-(D/E)XK superfamily endonuclease. The red bars indicate the regions detected by LIGR-Nue.

*Synechocystis* cells were incubated with AMT in microcentrifuge tubes and exposed to UV light at 365 nm to cross-link interacting RNA regions. Following rRNA depletion, S1 nuclease treatment was performed to remove non-crosslinked RNA. The cross-linked RNA duplexes were then subjected to proximity ligation using CircRNA ligase to generate chimeric RNA molecules before decrosslinking. RNA enrichment and decrosslinking were subsequently performed using RNase R treatment and UV irradiation at 264 nm, respectively. After reverse transcription, the resulting cDNA libraries were sequenced using an Illumina NovaSeq X Plus platform. Sequencing reads were mapped to the reference genome, and split reads representing the RNA–RNA interactions were identified using the RNAnue pipeline. Potential interactions were evaluated based on hybridization potential, sequence complementarity, and transcript abundance. The identified sRNA–mRNA interactions were subsequently visualized as a network using Cytoscape. A detailed stepwise experimental protocol is provided in **Supplementary Material 1**.

### Global RNA-RNA interactome of *Synechocystis*

A comprehensive analysis of the generated chimeric reads was performed to identify biologically relevant RRIs. The comparison in the number of interactions between AMT cross-linked (A+L+) and non-crosslinked (A-L+) samples in dark showed that overall chimeric interactions increased due to AMT cross-linking supporting the effectiveness of AMT crosslinking in capturing physiologically relevant RRIs (**Supplementary Figure S2 A**). Notably, 8.48-fold increase in sRNA–mRNA and 8.06-fold increase in sRNA-rRNA interactions were observed in A^+^L^+^ samples as compared to A^−^L^+^. The Circos plots of all the interactions generated from AMT crosslinked and proximity ligated samples revealed a broad genomic distribution of intra- and inter molecular RNA–RNA interactions involving multiple RNA transcript types (**Supplementary Figure S2 B-D**). Interaction arcs were distributed across the chromosome and naturally occurring plasmids, indicating that RRIs were not restricted to a small number of genomic regions. The interactions observed were consistent among all biological replicates, as depicted by the individual coverage tracks. To simplify, the fraction of each interaction type of the global RNA-RNA interactome is presented as pie chart in **Supplementary Figure S3**. The largest fraction among all the interactions consisted of rRNA–mRNA and mRNA-mRNA interactions. Interestingly, mRNA-mRNA interactions were just as prevalent in the dark as in PA and HL. We also observed a substantial number of mRNA-rRNA interactions in the PA and HL conditions, likely reflecting ongoing translation **(Supplementary Figure S3)**. There were less mRNA-rRNA interactions in dark, matching the shutdown of cyanobacterial metabolism in this condition **(Supplementary Figure S3)**. The pie-of-pie highlights remaining interaction classes, including mRNA–tRNA, rRNA–sRNA, sRNA–tRNA, mRNA–IS (mobile genetic element) interactions, etc.

As the primary focus of this study was to identify the RNA targets of sRNAs, the dataset generated for AMT-crosslinked and proximity-ligated samples under all experimental conditions was further filtered to extract the sRNA-associated interactions. Circos plot visualization revealed that sRNA-associated interactions were distributed across multiple regions of the *Synechocystis* genome (**Figure 4 A-C).** The inner tracks depicting the interaction-arm coverage derived from individual biological replicates showed consistent distribution of interaction-associated signals across the genome. **Figure 4 D-F** shows the subset circos plot generated for sRNA-mRNA interactions highlighted in red against the grey background of other RNA–RNA interaction classes. The coverage track represents the mean coverage derived from the biological replicates and highlights the enriched interacting regions on the genome. Interestingly, the number of sRNA-mRNA interactions increased significantly in dark as compared to the PA and HL (**Supplementary Figure S3 and Figure 4 D-F)**. Quantification of interaction classes further showed differences in both interaction abundance and contribution scores (**Figure 5**), providing a complementary measure of the relative representation of the different interaction classes. The average contribution score of these interactions was comparable among all three conditions; however, the number of interactions was relatively higher in the dark. This is consistent with the observed elevated total sRNA levels in the dark (**Figure 1**). Interestingly, the number of sRNA-rRNA interactions was comparable among all conditions, but they had relatively higher contribution score in dark as compared to PA and HL (**Figure 5**). This underlines the effectiveness of the experimental strategy in enriching sRNA–mRNA interactions by applying physiologically relevant stress conditions. Moreover, to characterize the detected RNA–RNA interactions, scores of different parameters were generated by LIGR-Nue as described in Methods. A complete list of the sRNA-RNA interactions detected by LIGR-Nue along with the all the parameters is provided in **Supplementary Table S1.**

**Figure 4.**
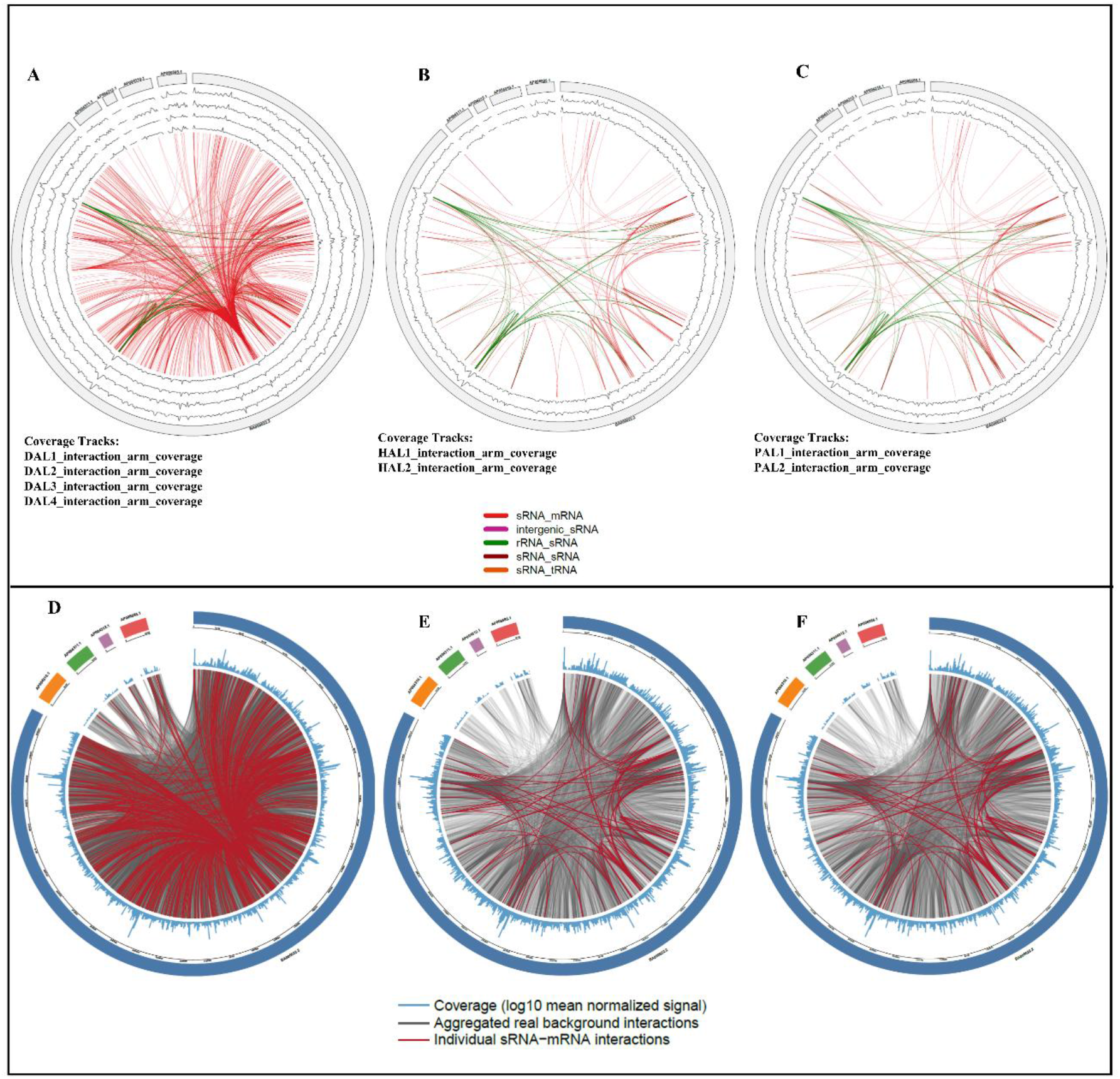
Circos Plots depict sRNA–RNA interactions across the *Synechocystis* genome under different conditions. (**A–C**) sRNA interactions with all other transcript types for which at least one interacting region was annotated as an sRNA. Interaction links or arcs are colored according to different transcript types, with sRNA–mRNA interactions highlighted in red under different conditions, Dark (A), HL (B) and PA (C). The outermost ring denotes the chromosome and four plasmids, with interaction-arm coverage represented by the inner tracks derived from the biological replicates. The arcs indicate the interacting RNAs at their corresponding positions. **(D-F)** Interaction map generated for sRNA-mRNA, indicated in red arcs, against all other interactions depicted in grey arcs under Dark (D), HL (E) and PA (F). The outer track represents the chromosome and four plasmids while the inner track represents the mean coverage derived from the biological replicates on log_10_ scale. For further details, see **Figure S2 and S3.**

**Figure 5.**
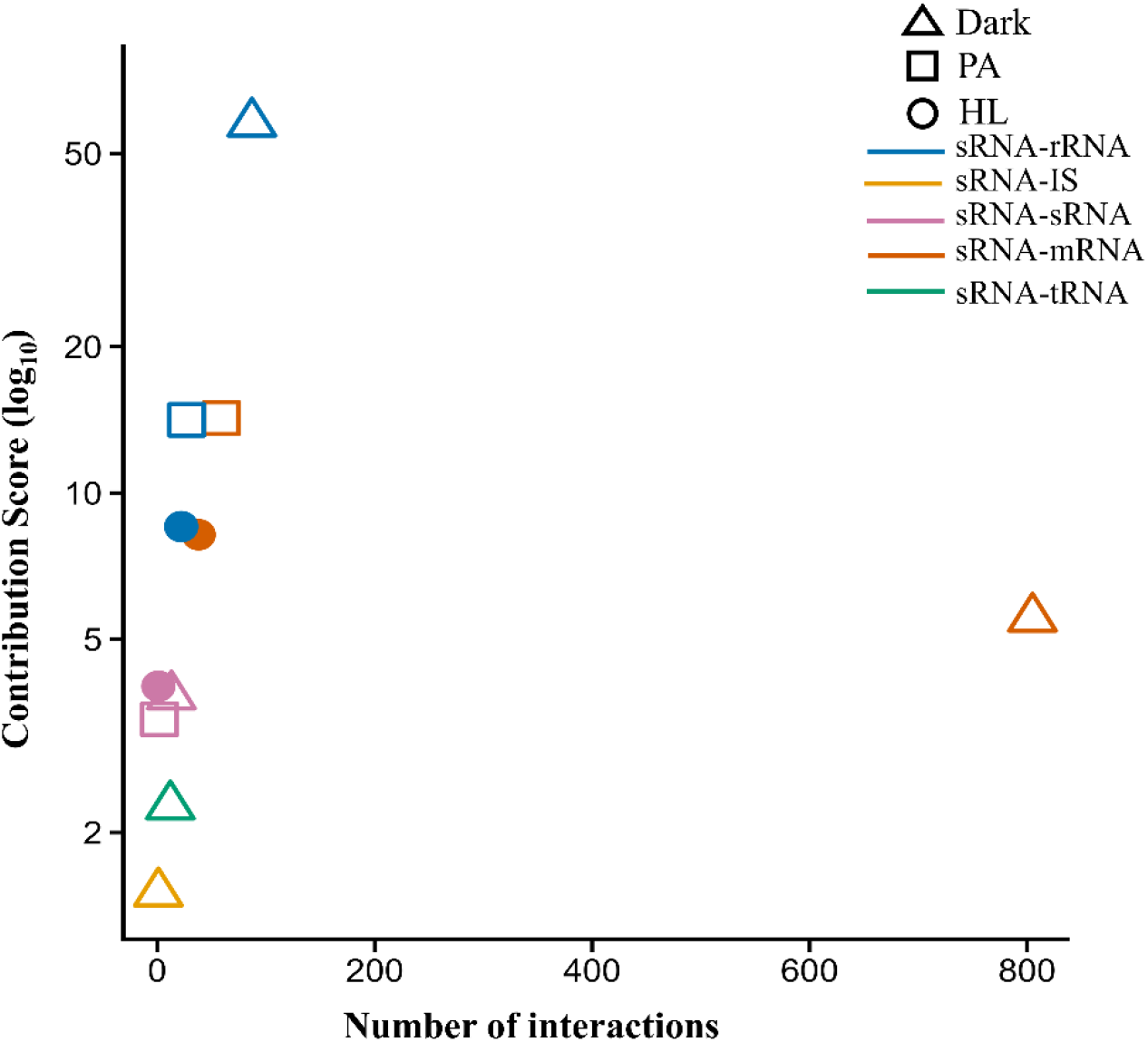
Distribution of sRNA-associated RNA–RNA interactions across transcript biotypes. The x-axis represents the number of detected interactions, and the y-axis represents the mean contribution score on a logarithmic scale. Symbols and colours represent experimental conditions .and interaction type respectively. Only sRNA associated interactions are shown. For further details, see **Figure S2 and S3**.

### Benchmarking against functionally characterized sRNAs

To validate the sRNA–mRNA interactions identified in this study, we compared our dataset with previously reported computationally predicted and experimentally validated targets, focusing on PsrR1 and NsiR4. PsrR1 is a post-transcriptional regulator of PSI–related genes during high-light stress. In *Synechocystis*, high-light stress strongly affects the photosynthetic apparatus, leading to a reduction in the number of photosystems and phycobilisome complexes. Therefore, a key strategy for high-light acclimation in cyanobacteria is the rapid downregulation of genes encoding photosynthetic pigments and photosystems, particularly PSI [51, 52]. Under high light, PsrR1 has been reported to target mainly the PSI-related genes *psaL, psaJ, psaI* and *psaF,* as well as other genes related to photosynthesis, namely *cpcA, psbB* and *chlN* [15, 18]. To determine whether LIGR-Nue detected previously reported PsrR1 interactions, we examined the targets identified in our dataset. Consistent with the earlier reports, interactions with *psaJ* and *cpcA* mRNAs were observed **(Figure 6A)**. In addition, interactions were observed with the *cpcC2* mRNA (gene *sll1579*) encoded just downstream of *cpcA,* and with the *ccmK1* and *rbcL* mRNAs **(Figure 6A)**. Under high-light conditions, the detected targets included *psaJ*, *rbcL cpcA, cpcC2,* and *ccmK1.* (**Supplementary Table S1**). To analyse further the base pairing regions between PsrR1 and its targets, the LIGR-Nue-detected chimeras were used as input for hybridization energy predictions by IntaRNA. The interaction between PsrR1 and *rbcL* was mapped to the 5′ untranslated region (5′-UTR), spanning the start codon (**Figure 6A**). This site has previously been reported as a bioinformatically predicted target [15]. Consistent with this observation, IntaRNA analysis predicted a stable interaction at the same site with a binding energy of −15.46 kcal/mol (**Figure 6A**).

**Figure 6.**
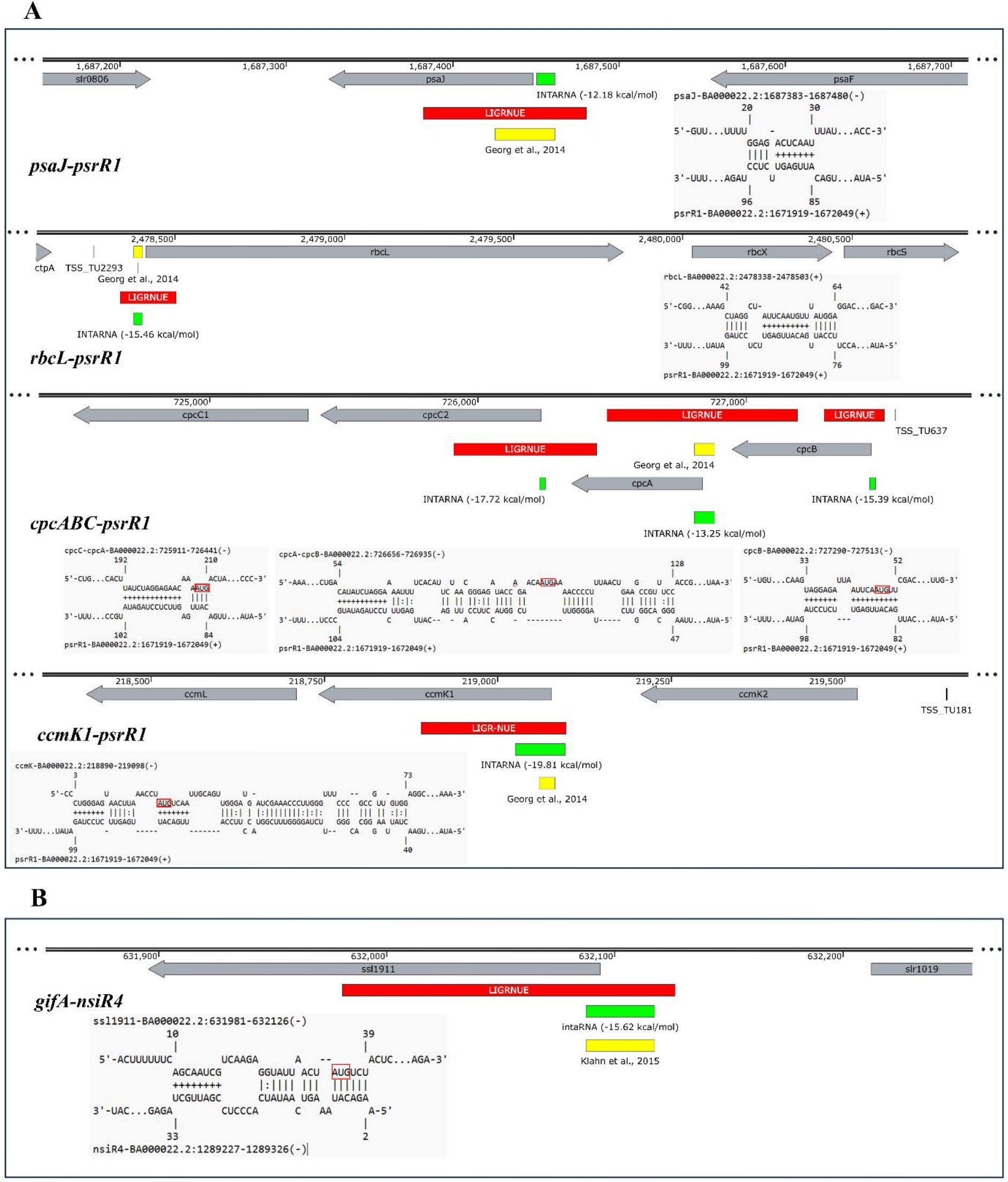
Benchmarking predicted sRNA–mRNA interactions against previously reported interactions. Snapshot of **(A)** PsrR1–target mRNA interaction regions showing predictions by LIGR-Nue (red) and IntaRNA (green; hybridization energies in brackets) alongside previously reported interaction regions [yellow, 15]. **(B)** NsiR4*-gifA* identified by LIGR-NUE (red), IntaRNA (green; energy in bracket) and previously validated [yellow, 21]. The alignments show the interacting nucleotides between the target mRNA (top) and the sRNA (query, bottom) as predicted by IntaRNA. The respective start codons are highlighted in red box. In the interaction representation, **+** denotes base pairs within the predicted putative seed region, a short region of favourable base pairing that contributes to the initiation and stability of the RNA–RNA interaction; **|** indicates intermolecular base pairing, while **−** represents unpaired or gapped positions.

Similarly, the binding sites of other PsrR1 targets were examined. The interaction site for *psaJ* was mapped to its 5′-UTR extending into the *psaJ* coding sequence (CDS), overlapping with the experimentally validated site. In agreement with this observation, IntaRNA predicted a binding site overlapping with the LIGR-Nue–detected region, with a hybridization energy of −12.18 kcal/mol. *cpcA* showed consistent interaction across all conditions. The binding region of *cpcA* spans a broad area encompassing part of the *cpcB* CDS, the 5′-UTR of *cpcA*, and the *cpcA* CDS, including the start codon. This region overlaps with both the experimentally validated binding site [15] and the IntaRNA-predicted site, with an interaction energy of −13.25 kcal/mol. In addition, LIGR-Nue detected a previously unreported binding site spanning the start codon of *cpcC2*, with an IntaRNA-predicted hybridization energy of −17.72 kcal/mol and another binding site at the start codon of *cpcB* with an IntaRNA-predicted energy of −15.39 kcal/mol **(Figure 6A)**. The binding site for *ccmK1* was mapped near its start codon, coinciding with both the IntaRNA prediction (−19.81 kcal/mol) and the previously reported interaction site **(Figure 6A)**. Interestingly, interaction of PsrR1 with *psaA* was also detected along with some other RNAs which are listed in **Supplementary Table S1 and S2**.

Another sRNA, NsiR4, has been reported to post-transcriptionally regulate the *gifA* mRNA, encoding the glutamine synthetase–inactivating factor IF7, under nitrogen-limiting conditions and an inverse relationship between NsiR4 abundance and IF7 accumulation has been shown [21]. This NsiR4-dependent regulation of *gifA* mRNA affects the cellular glutamine pool and consequently impacts NH₄⁺ assimilation through glutamine synthetase (GS) [21]. Consistent with this report, LIGR-Nue detected an interaction between NsiR4 and *gifA* near the start codon, spanning the 5′-UTR and CDS. IntaRNA analysis further predicted a strong interaction at the same region with a hybridization energy of −15.62 kcal/mol **(Figure 6B)**. In addition to *gifA*, several other mRNA targets—including *hisD, ccmK2, sll1697, sll0844* (tRNA-specific 2-thiouridylase MnmA)*, thrC* and an intergenic region between *ppc* (phosphoenolpyruvate carboxylase) and *slr0935* were identified, predominantly under the dark condition (**Supplementary Table S1**). A summary of all predicted PsrR1 and NsiR4 targets, along with their corresponding interaction scores, is provided in **Supplementary Table S1** and IntaRNA-derived hybridization energies in **Supplementary Table S2**.

### Genome-wide identification of sRNA–RNA interactions by LIGR-Nue

Following the confirmation of previously reported sRNA–mRNA interactions detected using LIGR-Nue, we further analysed the RNA targets of additional abundant sRNAs. LIGR-Nue interactions were retained in most cases, if detected in at least two biological replicates or under two distinct conditions, with IntaRNA-predicted interaction energies of less than −5 kcal/mol. A complete list of interactions meeting these criteria is provided in **Supplementary Table S1**. Interactions with higher split read counts (no_splits) represented interactions detected more frequently across the sequencing data, while high complementarity (GCS) and hybridization scores (GHS) suggested favourable base-pairing and stable binding. Some interactions showed strong GCS and GHS despite low crosslink values, likely reflecting discontinuous binding patterns typical of sRNA interactions. Together, these parameters enabled identification of biologically relevant interactions across conditions. The corresponding IntaRNA-predicted structures and hybridization energies are shown in **Supplementary Table S2 and Supplementary Material 2**. In addition, to better visualise the interactions detected, sRNA-RNA interactions ≤ −10 kcal/mol hybridization energies were considered to build the interaction network **(Figure 7)**. Notably, the majority of the interactions were consistently detected across multiple conditions with overlapping binding regions on the same target transcripts. The recurrence of these interactions at similar target sites across independent samples suggests that the identified sRNA–RNA interactions represent specific and reproducible binding events rather than random ligation artifacts.

**Figure 7.**
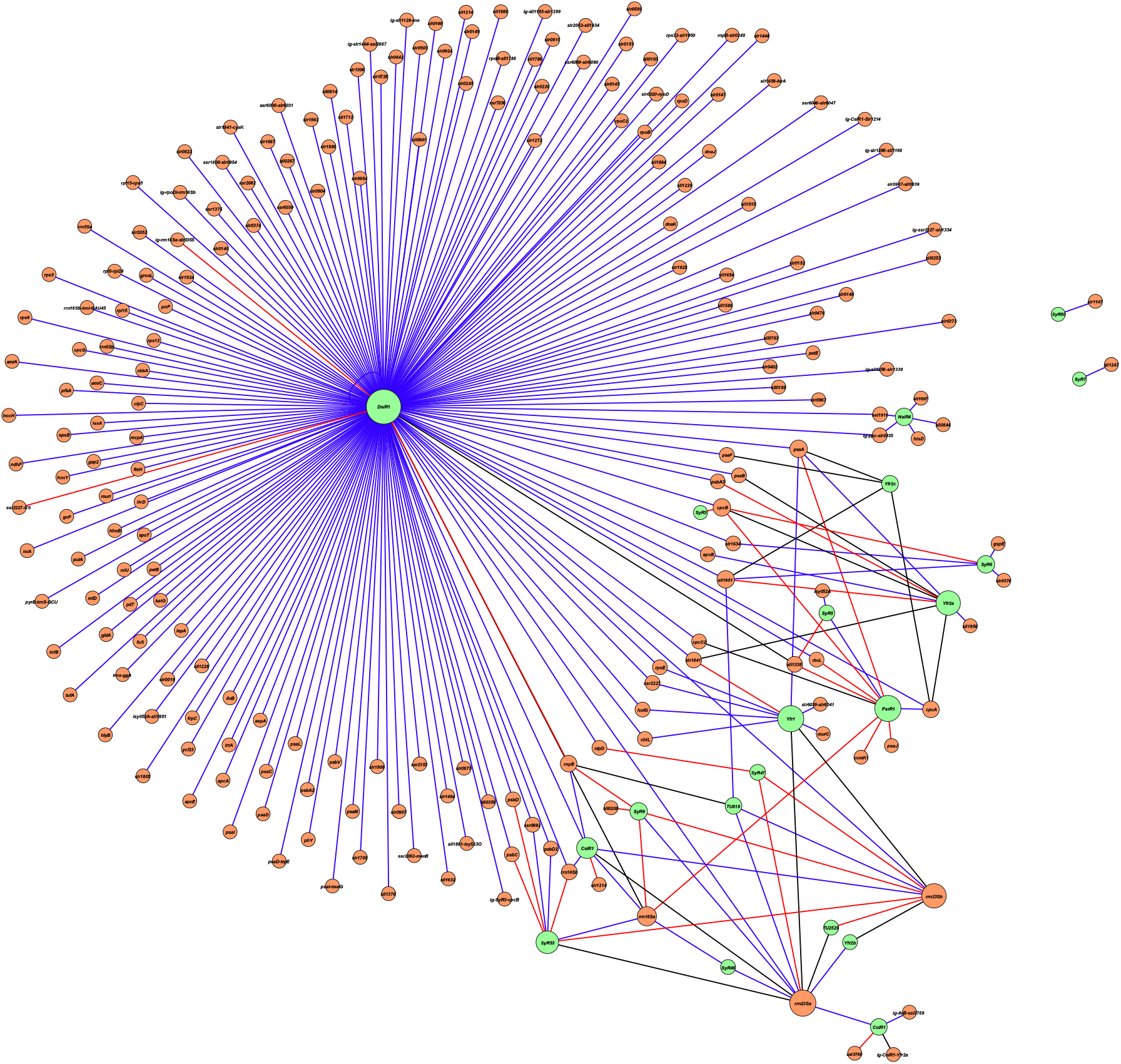
Network of sRNA–target RNA interactions in *Synechocystis*. sRNA-RNA interactions ≤ −10 kcal / mol hybridization energies were considered to build the interaction network. Green nodes represent sRNAs and orange nodes represent their target RNAs. Node size is proportional to degree, reflecting the number of connections associated with each node. Edges denote predicted interactions, with their widths scaled according to IntaRNA interaction energy to indicate the relative strength of predicted interactions and their colours indicating the condition: PA (black), HL (red), and dark (blue). If the interacting region spans two consecutive genes, the names of both genes are included. Interactions mapped to intergenic regions are prefixed with “ig”. DsiR1 exhibits the highest number of interactions under dark conditions consistent with its abundant accumulation under dark.

Regulatory sRNAs are sometimes located adjacent to the transcriptional units regulated by them. According to the LIGR-Nue results, at least 8 abundant sRNAs interact with transcripts of genes located adjacent to them **(Table 1)**. SyR3 and its target *cpcB* are oriented away from each other, resembling an archetypical arrangement. However, in all other instances like DsiR1/*ssr2227*, CsiR1/*slr1214*, NsiR4/*sll1697* and SyR53/*psbC* & *psbD*, the sRNAs and the identified mRNAs belong to the same transcriptional unit. This points to the likely underestimated role of 3’- and 5’-UTR-derived sRNAs, a recently recognized source of regulatory RNAs [53, 54].

**Table 1:**
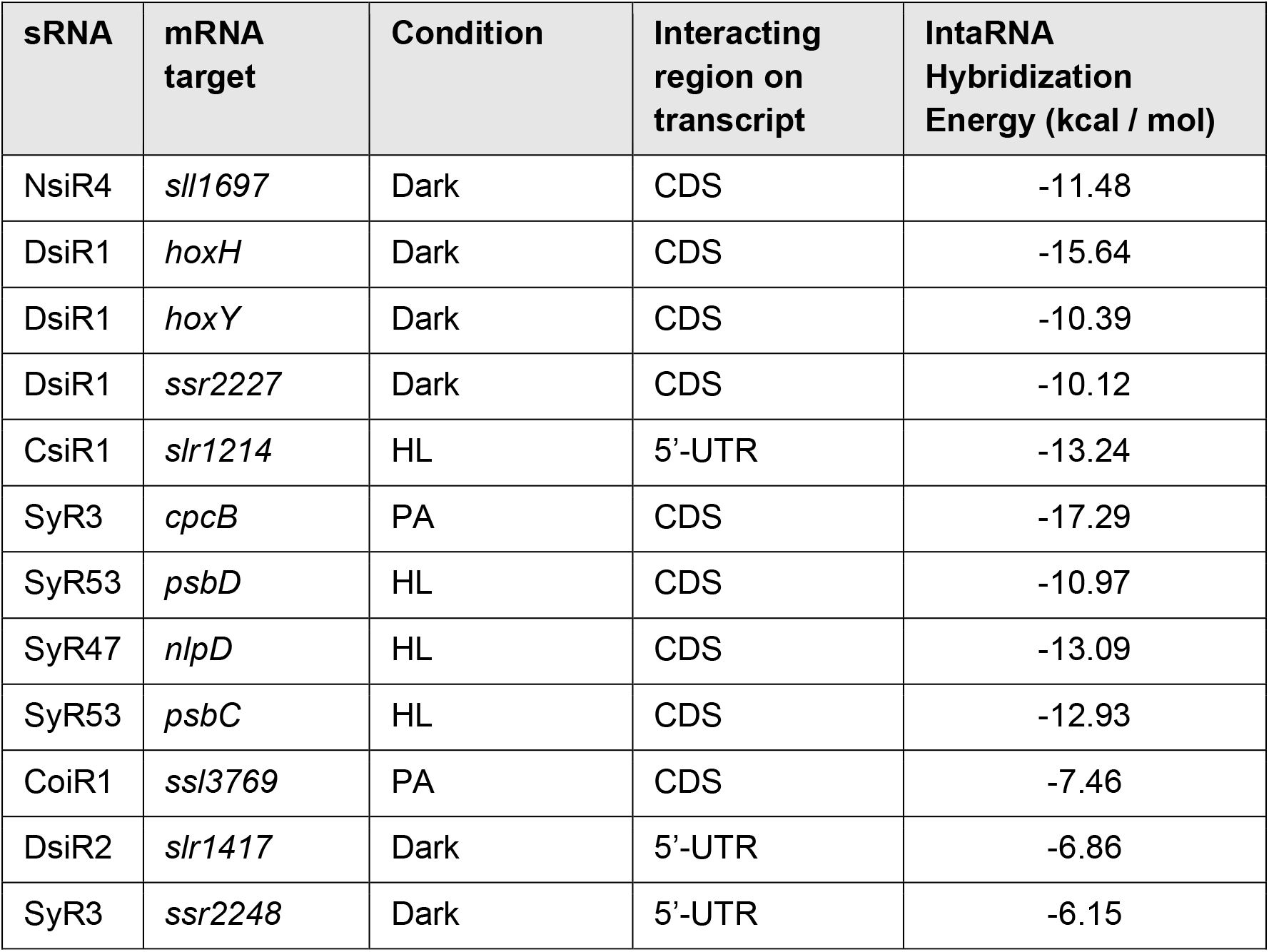
List of sRNAs located adjacent to their target mRNAs in the *Synechocystis* genome.

| **sRNA** | **mRNA target** | **Condition** | **Interacting region on transcript** | **IntaRNA Hybridization Energy (kcal / mol)** |
|---|---|---|---|---|
| NsiR4 | *sll1697* | Dark | CDS | -11.48 |
| DsiR1 | *hoxH* | Dark | CDS | -15.64 |
| DsiR1 | *hoxY* | Dark | CDS | -10.39 |
| DsiR1 | *ssr2227* | Dark | CDS | -10.12 |
| CsiR1 | *slr1214* | HL | 5’-UTR | -13.24 |
| SyR3 | *cpcB* | PA | CDS | -17.29 |
| SyR53 | *psbD* | HL | CDS | -10.97 |
| SyR47 | *nlpD* | HL | CDS | -13.09 |
| SyR53 | *psbC* | HL | CDS | -12.93 |
| CoiR1 | *ssl3769* | PA | CDS | -7.46 |
| DsiR2 | *slr1417* | Dark | 5’-UTR | -6.86 |
| SyR3 | *ssr2248* | Dark | 5’-UTR | -6.15 |

IntaRNA-predicted base-pairing regions were identified for these sRNA–mRNA pairs either in the 5′-UTR or within the CDS. In addition to these cis interactions, LIGR-Nue also detected separate targets for these sRNAs, supported by significant LIGR-Nue interaction scores and IntaRNA-predicted interactions with favourable hybridization energies (**Supplementary Table S1 and S2, Supplementary Material 2**).

DsiR1 is the most abundant sRNA in *Synechocystis*, especially under darkness (**Figure 1**). Consistent with its expression, LIGR-Nue identified numerous RNA interactions associated with DsiR1, predominantly in the dark condition (**Figure 7**). The hybridization energies of these interactions were subsequently predicted using IntaRNA. The detected mRNA targets included genes encoding photosystem and phycobilisome components like *psaA, psaB, psaC, psaD, psaI, psaF, psaL, psaM, psbC, psbD, psbA2, psbA3, psbD2, psbV, apcA, apcB, apcE, cpcA, cpcB, cpcC,* and *cpcG* (**Figure 7 and Supplementary Table 1**). On the majority of these highly abundant transcripts, DsiR1 base-pairs at multiple sites. In addition, IntaRNA analysis revealed DsiR1 interacts at specific sites on these transcripts with significant hybridization energies (**Supplementary Table 2 and Supplementary Material 2**). These highly expressed genes are known to be rapidly downregulated during light to dark transition. It is highly likely that the rapidly up-regulated levels of DsiR1 under dark trigger rapid degradation of these transcripts, thereby contributing to downregulation of these photosynthetic genes under dark. DsiR1 was also observed to interact with mRNAs encoding the core hydrogenase components (*hoxH* and *hoxY*) and putative transposase *ssr2227* (**Supplementary Figure S4**), as well as alternative sigma factors *rpoB, rpoD* and *rpoE* along with core subunits of RNA polymerase *rpoC1* and *rpoC2* **(Supplementary Table S1, S2 and Supplementary Material 2).** DsiR1 also base-pairs with transcripts of several transcripts such as *slr1924, lexA, slr1306, slr1805 (hik16), slr0975, murI*, *clpC, sll1338, slr0476, gcp, rbcL* and *slr1474, slr0249, slr1841, dnaK, hofG, frpC, slr0442, sll1951,* and *ssr7036*, with significant hybridization energies. Most of these targets are down-regulated under darkness. The *slr0144-slr0151* operon contains overlapping structural genes encoding a vinyl reductase, or accessory proteins during the assembly of the PSII complex. LIGR-Nue detected interactions of DsiR1 with *slr0146*, *slr0147, slr0149* and *slr0151* within this operon. In several cases, DsiR1 was found to interact within the coding regions of target mRNAs, apart from the 5′-UTRs.

Another sRNA, CsiR1, which strongly accumulates under dark stress (**Figure 1**) and is originating directly upstream of *slr1214*, was found to interact with the *slr1214* mRNA with a significant hybridization energy (**Supplementary Figure S5 and Table 1**).

Interestingly, in addition to sRNA–mRNA interactions, LIGR-Nue also detected interactions of several sRNAs—including DsiR1, CoiR1, CsiR1, PsrR1, SyR5, SyR9, SyR46, SyR47, SyR53, TU2529, TU818, Yfr1, and Yfr2b—with specific regions of the ribosomal RNAs *rrn23S* and *rrn16S* (**Supplementary Figure S6 and Figure 7).** Notably, DsiR1 showed base-pairing with multiple regions across the 23S, 16S, and 5S rRNAs in the dark condition, and with mRNAs encoding the translation-related proteins InfB, Rps5 and Rpl15 with significant hybridization energies as predicted by IntaRNA. These observations suggest a potential impact of certain sRNAs on the translational machinery, although the functional relevance of these interactions remains to be investigated (**Supplementary Figure S6 and Supplementary Material 2**).

## DISCUSSION

In this study, we present LIGR-Nue **(Figure 2 and Supplementary Material 1)** as an *in vivo* strategy for generating global RRI maps in the cyanobacterium *Synechocystis*. To our knowledge, this represents the first global RNA interactome analysis performed in a cyanobacterial system. LIGR-Nue complements existing approaches for probing RNA duplexes in cyanobacteria by enabling direct detection of RNA–RNA interactions within intact cells under physiologically relevant conditions. Conceptually analogous to proximity ligation methods used to map long-range chromatin interactions in humans [32], this approach captures both canonical and non-canonical RNA interactions that are inaccessible to *in vitro* methods, where RNA folding and pairing occur outside the native cellular context. *In vivo* RNA duplex crosslinking using AMT preserves interacting RNA regions from degradation by S1 nuclease. A critical step in this method is ribodepletion following crosslinking, as rRNA constitutes approximately 90% of the total cellular RNA. Despite ribodepletion, our dataset still revealed both intra- and intermolecular rRNA interactions. In addition, duplex RNA species in which rRNAs base-pair with other transcript types were also detected. Therefore, the presence of a substantial fraction of rRNA-associated interactions in the dataset is in agreement with the fact that rRNA represents the predominant RNA class in the cell (**Supplementary Figure S3**). Another important strategy was to subject cells to abiotic stress conditions to enhance the abundance of specific sRNAs. We specifically selected darkness, as the sRNA DsiR1, along with several others, showed strong accumulation under this condition (**Figure 1**). In addition, high-light stress was included to allow cross-validation of LIGR-Nue–derived interactions with previously reported sRNA– mRNA interactions. Enriching sRNAs under diverse stress conditions therefore represents an important initial step toward capturing physiologically relevant *in vivo* sRNA–mRNA interactions and enabling their subsequent detection and analysis.

The robustness of the LIGR-Nue method is further strengthened by integration with the RNAnue analysis pipeline (https://github.com/RNABioInfo/RNAnue/releases/tag/v1.0.0-beta.1) [33], which is specifically designed to infer RNA–RNA interactions from duplex-directed sequencing data. Unlike conventional RNA-seq–based workflows, RNAnue incorporates hybridization potential, thermodynamic stability, and rigorous statistical assessment to filter chimeric reads, thereby substantially improving both the specificity and reliability of interaction predictions. In the un-crosslinked and non-ligated samples, RNAnue detected a chimeric RNA mapped to two genomic locations, derived from exon fusion of tRNA_i_^fmet^ (**Figure 3**). This chimera was not observed in the AMT crosslinked and proximity-ligated samples, as it originated from exon fusion rather than RNA–RNA hybridization. The detection of this intron-derived fragment (split read) in all non-ligated samples validated the specificity of the chimera detection approach implemented in the RNAnue bioinformatics pipeline. Furthermore, the detection of a relatively higher number of split reads in AMT-crosslinked samples provided additional evidence supporting successful AMT-mediated RNA crosslinking in *Synechocystis* cells (**Supplementary Figure S2A**). The predominance of low-to-moderate sRNA–mRNA interaction scores across the conditions is consistent with established models in which sRNAs regulate gene expression through multiple weak or transient interactions rather than strong binding to a few targets [55]. The increased interaction density observed under the dark condition suggests condition-dependent enhancement of sRNA-mRNA interactions commonly associated with adaptive regulatory responses (**Figure 4 & 5**). Especially in this study, DsiR1 strongly accumulated under dark and also exhibited the highest number of interactions under this condition (**Figure 1 & 7**).

Benchmarking against well-characterized sRNA–mRNA interactions demonstrated that LIGR-Nue reliably captured functionally relevant RNA–RNA interactions *in vivo* (**Figure 6**). Importantly, enrichment of sRNAs under different abiotic stress conditions relevant to photosynthetic cyanobacteria substantially increased their cellular abundance, enabling robust detection of condition-specific and biologically meaningful interactions that would otherwise remain undetectable under standard growth conditions. In this regard, the detection of already reported PsrR1 interaction sites under high-light conditions by LIGR-Nue is consistent with the proposed feed-forward regulatory loop involving the transcriptional regulator RpaB and the sRNA PsrR1. Upon a shift to high light, dual repression of PSI genes by RpaB at the transcriptional level and by PsrR1 at the post-transcriptional level is thought to provide a rapid and efficient means to suppress PSI gene expression by inhibiting translation and promoting accelerated degradation of the target mRNAs, thereby enabling rapid acclimatization to high light [15, 18]. The PsrR1 targets identified in this study (**Figure 6A**)—*rbcL*, *psaJ*, *cpcA*, *ccmK1*—encode functionally interconnected components of the photosynthetic and carbon assimilation machinery and were previously reported as PsrR1 targets [15]. Targeting *rbcL* likely affects translation of the entire operon because it is the first gene in the *rbcLXS* operon. The gene *rbcL* and *rbcS* encode the large and small subunits of Rubisco, the major Calvin Cycle enzyme for CO_2_ fixation, while *ccmK1* encodes a major protein of the carboxysome, a microcompartment that encapsulates Rubisco to boost carbon fixation. Therefore, these results point at PsrR1 as a regulator of carbon assimilation. In addition to known interactions of PsrR1 with *cpcAB* transcripts, LIGR-Nue also detected an interaction with *cpcC2.* Cpc2 is a phycobilisome rod linker polypeptide binding the phycocyanin α and β subunits. Therefore, its regulation in concert with *cpcA* and *cpcB,* which encode these subunits, is physiologically meaningful. The NsiR4–*gifA* interaction detected by LIGR-Nue is consistent with the previous report [21] (**Figure 6B**). The concordance between LIGR-Nue–detected interaction regions, previously validated binding sites, and IntaRNA predictions reinforces the robustness of our approach. In a previous proteogenomic analysis, several of the proteins encoded by these mRNAs (CpcB, PsaL, CpcA, CpcC2, PsaF, PsaA, CcmK1 and RbcL) were found to belong to the 20 most-abundant proteins in *Synechocystis* [56]. Therefore, tight control of their translation by interacting sRNAs is likely of high relevance.

The global RNA–RNA interaction landscape captured by LIGR-NUE provides additional insight into the roles of highly abundant and condition-responsive sRNAs during abiotic stress (**Supplementary Table 1 and Figure 7**). The comprehensive catalogue of sRNA– mRNA interactions identified in this study serves as a platform for characterizing their regulatory roles under a given abiotic stress condition. Out of the 33 abundant sRNAs identified, 8 were found to interact with mRNAs transcribed from adjacent genomic loci (**Table 1**). Notably, the pronounced accumulation of DsiR1 under dark conditions correlated with the large number of interactions detected by LIGR-Nue (**Figure 7**). Many of the identified targets encode components involved in photosynthesis, carbon metabolism, and redox processes, which are known to be downregulated or reprogrammed in the absence of light, consistent with a coordinated dark-acclimation response. Transcripts encoding PSI, PSII, and phycobilisome components accumulate at high levels under light conditions, but are rapidly degraded in the dark. Given their high cellular abundance, elevated levels of DsiR1 may facilitate their rapid turnover during the acclimation to darkness, potentially including a block on the translational apparatus via interactions with rRNAs. Moreover, interactions with global regulatory factors, including sigma factors and translation initiation components, suggest a role for DsiR1 in large-scale transcriptional and translational reprogramming. The exceptionally high abundance under dark conditions suggests that DsiR1 may act as a global post-transcriptional regulator facilitating metabolic and translational downshifting during prolonged darkness. Together, the functionally coherent set of targets identified by LIGR-Nue supports a central regulatory role for DsiR1 during energy limitation. Similarly, LIGR-Nue detected condition-dependent interactions involving CsiR1, especially under dark, that are consistent with its previously proposed role in carbon stress responses [16]. In particular, the interaction of CsiR1 with *slr1214*, encoding the PatA-type phototaxis regulator NixC/LsiR [57, 58], located downstream of CsiR1, suggests a potential role in stress acclimation.

A known limitation of LIGR-seq is that the distances between ligation junctions and underlying RNA duplexes can vary substantially, as local secondary and tertiary RNA structures restrict S1 nuclease accessibility prior to ligation [32]. In line with this observation, we frequently observed extended interaction regions in which a single sRNA appeared to contact large portions, or in some cases nearly the entire open reading frame of a target mRNA. One plausible explanation is that highly abundant sRNAs form stable, multi-contact interaction interfaces with their targets, which protect extended RNA regions from nuclease digestion and result in longer ligation products.

In conclusion, LIGR-Nue reliably captured physiologically relevant sRNA–mRNA interactions in *Synechocystis*, with detected targets aligning with the known sRNA functions, expression patterns, and stress-specific abundance. The strategic use of the stress conditions proved effective in enriching condition-responsive sRNAs and uncovering meaningful regulatory interactions that would otherwise remain undetected. Together, these results validate LIGR-Nue as a robust platform for mapping RNA–RNA interaction networks in bacteria and provide a systems-level framework for investigating the regulatory models. By preserving real-time RNA interactions influenced by cellular physiology and environmental cues, LIGR-Nue offers a systems-level perspective on the post-transcriptional regulatory networks of *Synechocystis*. Beyond method development, the resulting interaction dataset represents a versatile resource that can be mined for sRNA-specific or condition-specific hypotheses and serves as a foundation for targeted experimental validation and future mechanistic studies of post-transcriptional regulation in photosynthetic bacteria.

## ACKNOWLEDGEMENTS

Library preparation and sequencing was performed by Chromosome labs Pvt. Ltd (Mumbai, India). We thank Dr. Ashwani Kumar, Application expert, New England Biolabs, for the support.

## AUTHOR CONTRIBUTIONS

Conceptualization and Supervision: J.S.S.P. Methodology, Investigation and Writing-Original Draft: M. M. Bioinformatics analysis: C.A. and J.G. Formal analysis: J.S.S.P, W.R.H., B.V., S.K. Writing-review & editing: J.S.S.P, W.R.H., B.V. Resources: J.S.S.P, W.R.H, B.V. Funding: W.R.H. B.V and J.S.S.P.

## SUPPLEMENTARY DATA

Supplementary Data are available at NAR online.

## DATA AVAILABILITY

RNA-Seq datasets were deposited in the Gene Expression Omnibus (GEO) database, with accession number GSE353050.

(direct access:https://www.ncbi.nlm.nih.gov/geo/query/acc.cgi?acc=GSE353050)

