## Supplementary material 1 and supplementary figures for "LIGR-Nue for the global mapping of the bacterial sRNA targetome *in vivo*"

**Supplementary Material 1: LIGR-Nue Protocol**

**Reagents**

- 4′-Aminomethyltrioxsalen hydrochloride (AMT) (Sigma- A4330)
- RNase-free phosphate-buffered saline (PBS), pH 7.4 (Thermo Fisher, AM9624)
- TS buffer (Tris-HCl and NaCl pH: 7.5)
- ATP (100 mM, Thermo fisher, R0441)

**Enzymes**

- TURBO DNase (Invitrogen, AM1907)
- Proteinase K (RNase-free, Millipore, 70663-4)
- S1 nuclease (Thermo fisher, EN0321)
- CircLigase ssDNA ligase (Epicentre, CL4111K)
- RNase R (Epicentre, RNR07250)

**Ribodepletion and RNA purification kit**

- NEBNext® rRNA Depletion Kit (Bacteria) with RNA Sample Purification Beads (E7860)
- Qubit™ RNA High Sensitivity (HS) (Invitrogen, Q32852)

**Reagent setup**

1. Prepare 5x TS buffer (100 mM Tris-HCl pH 7.5, 1.5 M NaCl).
2. Prepare AMT (1 mg/mL) in RNase-free water and keep on ice in dark.

**AMT crosslinking setup**

1. Arrange the UV crosslinking system using three 365 nm UV tube lights (TL-D 15W Actinic BL UVA A3, Philips) mounted in parallel close to each other at a distance of 7 cm as shown in Figure 2.
2. Position the glass petri-plates at a uniform distance of ~15 cm from the UV source to ensure consistent irradiation across all plates (9 cm diameter).
3. On the day of crosslinking, prepare the AMT solution containing 1x TS buffer and 0.3 mg $mL^{-1}$ AMT in nuclease-free water.
4. Collect the cells (~$2.5 \times 10^9$ cells) from culture tube and centrifuge at 2100 x g for 5 min at 34 °C, then discard the supernatant.
5. Wash pellet twice with 500 µL cold PBS and finally with 1 x TS buffer. After each wash centrifuge at 18000 x g for 1 minute and discard the supernatant.
6. Suspend cell pellet in 1 ml AMT solution (for samples to be crosslinked) and 1x TS (for control samples) and incubate on ice for 15 min in low light, mixing every 1 min.
7. Spread the cell suspension evenly onto plate (1 ml per plate) to form a uniform single layer, ensuring consistent irradiation of all cells. Maintain the sample on ice throughout the irradiation period.
8. Irradiate using 365 nm UV, 15 mW / $cm^2$ for 10 min × 6 cycles. Mix well between the irradiation phases as shown in Figure 2.
9. Collect cells and store pellet at −80 °C or proceed for the subsequent steps.

**RNA isolation**

RNA isolation was done as described in Los et al., 1997. RNA concentration was estimated by nanodrop and Qubit fluorometer using Qubit RNA HS kit (Invitrogen, Q32852) according to manufacturer's manual.

**Ribosomal RNA depletion by NEBNext® rRNA depletion kit (bacteria) with RNA sample purification beads (NEB, E7860)**

1. Anticipating RNA loss during subsequent steps, scale up the reaction to ensure sufficient RNA yield for downstream sequencing. The concentration of the input RNA is 4 µg. Make the following reaction mixture:

| Probe hybridization reaction | Volume (µl) |
|---|---|
| Total RNA in nuclease-free water (4 µg) | 44 |
| NEBNext Bacterial rRNA depletion solution | 8 |
| NEBNext Probe hybridization buffer | 8 |
| Total | 60 |

2. Split the reaction mixture into two PCR tubes to accommodate thermocycler volume requirements.
3. Mix the reaction components thoroughly by pipetting and spin briefly. Place the tubes in the thermocycler. Set thermocycler with following parameters and lid temperature 105 °C:

| Temperature | Time |
|---|---|
| 95 °C | 2 min |
| Ramp to 22 °C | 0.1 °C/s |
| Hold | 5 min |

4. Run the program. After the run, immediately put the tubes on ice.

5. Proceed to RNase H digestion. Into these tubes, add the following components:

| RNase H digestion reaction | Volume per tube (µl) |
|---|---|
| Hybridized RNA from previous reaction | 30 |
| RNase H reaction buffer | 4 |
| NEBNext Thermostable RNase H (5 U/µL) | 4 |
| Nuclease-free water | 2 |
| Total volume | 40 |

6. Mix thoroughly and incubate at 50 °C for 30 min in the thermocycler with 55 °C lid temperature.
7. Briefly spin down the tube in a microcentrifuge and place on ice. Proceed immediately to DNase I digestion
8. Assemble the following DNase I digestion reaction on ice (pool the reactions into 1.5 ml microfuge tube).

| DNase I master mix | Volume (µl) |
|---|---|
| RNase H treated RNA | 80 |
| DNase I reaction buffer | 20 |
| NEBNext DNase I (RNase-free) (20 U/µL) | 10 |
| Nuclease free water | 90 |
| Total volume | 200 |

9. Mix thoroughly by pipetting and briefly spin down the tube in a microcentrifuge. Incubate in a heating block for 30 minutes at 37°C. Briefly spin down the tube in a microcentrifuge and place on ice. Proceed immediately to RNA purification.

**RNA purification using NEBNext RNA sample purification beads**

1. Vortex gently the NEBNext RNA sample purification beads to resuspend for homogenized mixture.
2. Collect the samples in 2 ml micro-centrifuge tubes and add 360 µl (1.8x for 200 µl reaction volume) beads to RNA sample from previous step and mix thoroughly by

pipetting up and down. Incubate for 15-20 min on ice to allow binding of RNA to the beads
3. Place the tubes on magnetic rack to separate the beads from the supernatant. After the solution is clear, carefully remove and discard the supernatant. Be careful not to disturb the beads which contain the RNA.
4. Wash twice with 800 µl of freshly prepared 80% ethanol to the tube while in the magnetic rack.
5. Completely remove residual ethanol and air dry the beads at room temperature for up to 5 min while the tube is on the magnetic rack with the lid open. Caution: Do not over-dry the beads. This may result in lower recovery of RNA target. Elute the samples when the beads are still dark brown and glossy looking, but when all visible liquid has evaporated. When the beads turn lighter brown and start to crack, they are too dry. Do not use thermostat for drying.
6. Remove the tube from the magnetic rack. Elute the RNA from the beads by adding 26 µl of nuclease-free water. Mix thoroughly by pipetting up and down at least 10 times and briefly spin the tube. Incubate for 5 minutes at room temperature.
7. Place the tube on the magnetic rack until the solution is clear (~ 5 minutes). Collect the supernatant containing RNA and transfer to a nuclease-free tube (1.5ml). Collect as much as possible without disturbing the beads. Place the tube on ice and proceed with downstream application. Alternatively, the sample can be stored at -80°C.
8. Measure the concentration of RNA. Ideally, ~200-300 ng RNA will be obtained. Use entire RNA for S1 nuclease digestion.

**S1 Nuclease digestion:**

1. Set up the S1 nuclease digestion reaction
   Step 1: Dilute 5x S1 nuclease buffer to 1x (add 20 µl of buffer to 80 µl of nuclease free water)
   Step 2: In a separate tube add 1µl of S1 nuclease (100 U) to 99 µl of S1 nuclease 1x buffer)
2. Prepare the following reaction:

| **Reagents** | **Volume (µl)** |
|---|---|
| S1 nuclease buffer (5x) | 8 |
| RNA (depends on concentration) | x |
| S1 nuclease<br>(1:100 dilution in 1x S1 nuclease buffer) – add from step 2 | 4 |
| Total | 40 |

3. Incubate the mixture at room temperature for 30 mins.
4. Proceed for bead-based purification. As the reaction volume is 40 µl, 1.8x beads to be added (72 µl) and mixed thoroughly. Incubate for 20 min on ice.
5. Keep the vials on magnetic stand and allow for solution to become clear. Discard the supernatant. Wash with 150 µl 80% ethanol and repeat the step. The amount of ethanol depends on the fact that it should submerge the beads. Remove residual ethanol and air dry at RT. Elute in 7 µl RNAse free water.

**Proximity ligation with CircRNA Ligase:**

1. Set the thermocycler program:
   85 ºC: 2 min
   60 ºC: 1 h

2. Add 2µl 10x CircRNA ligase buffer and nuclease free water to entire S1 digested RNA and keep in thermocycler for 2 min at 85 ºC to denature RNA (Note: This step is for all the samples).

3. Immediately place the tubes on ice to prevent renaturation and add 1µl of 10 mM ATP, 1µl $MnCl_2$. Add 1 µl of CircRNA ligase (100 U/µL) only for samples to be proximally ligated and add nuclease free water to the rest of the samples. Total reaction volume is 20 µl. Incubate the reaction at 60 °C for 1 h in thermocycler.

**Enrichment of cross-linked RNA and crosslink reversal**

1. Set the following reaction:

| **Reagents** | **Volume (µl)** |
|---|---|
| RNase R (20 U/µL) | 0.5 |
| 10x Reaction Buffer | 2.5 |
| Sterile water | 2 |
| Ligation reaction mixture from previous step | 20 |
| Total | 25 |

2. Mix the reaction components properly and incubate for 10 min at 37 °C.
6. Immediately after incubation, place the tubes on ice and extract RNA using bead-based purification. Use 45 µl beads (1.8 x for 25 µl reaction vol).
7. Wash twice with 200µl 80% ethanol and resuspend the RNA in 20 µl of 1x TS buffer in 2 ml micro-centrifuge tube.
8. For crosslink reversal, place the 2 ml microfuge tubes on ice and keep the caps open.
9. Irradiate the RNA for 5 min twice at 254 nm at 15 cm from three UV bulbs (TUV 15W G15/T8 L2, Philips) mounted in parallel with an interval of 5 min. Keep on ice during irradiation phases.
10. Remove AMT by ethanol precipitation and resuspend RNA in 10 µl of TS buffer.
11. Proceed for library preparation and deep sequencing. (Details in Methods)

**Supplementary Material 2:** RNA–RNA hybridization structures predicted by IntaRNA for the interaction pairs identified by LIGR-Nue. Structures are provided for the corresponding sRNA–target RNA pairs. See separate Text File

**Supplementary Material 3:** Annotation File used by LIGRNue. See separate .txt file

**Supplementary Table S1:** Complete catalogue of sRNA–target RNA interactions identified by LIGR-Nue across different experimental conditions. Each sheet corresponds to individual sRNA and labelled accordingly. See separate Excel File

**Supplementary Table S2:** Predicted hybridization energy scores, calculated using IntaRNA, for RNA–RNA interaction pairs identified by LIGR-Nue. Individual worksheets correspond to each sRNA, with scores arranged in ascending order. Lower energy values indicate more favourable predicted hybridization. See separate Excel File.

**Supplementary Figures:**

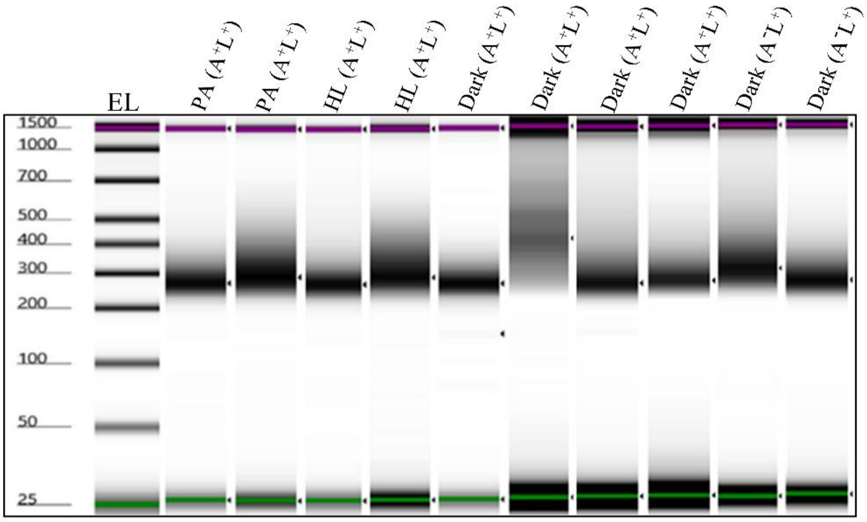


**Figure S1:** Tape Station profiles of the LIGR-Nue libraries. EL denotes electronic ladder with fragment sizes indicated in base pairs. PA, Photoautotrophic; D, dark stress (1 h); HL-High light stress (30 min). $A^+$ denotes AMT crossinking, while $A^-$ denotes samples not crosslinked with AMT. $L^+$ , proximity ligation with CircLigase; $L^-$ , non-ligated samples.

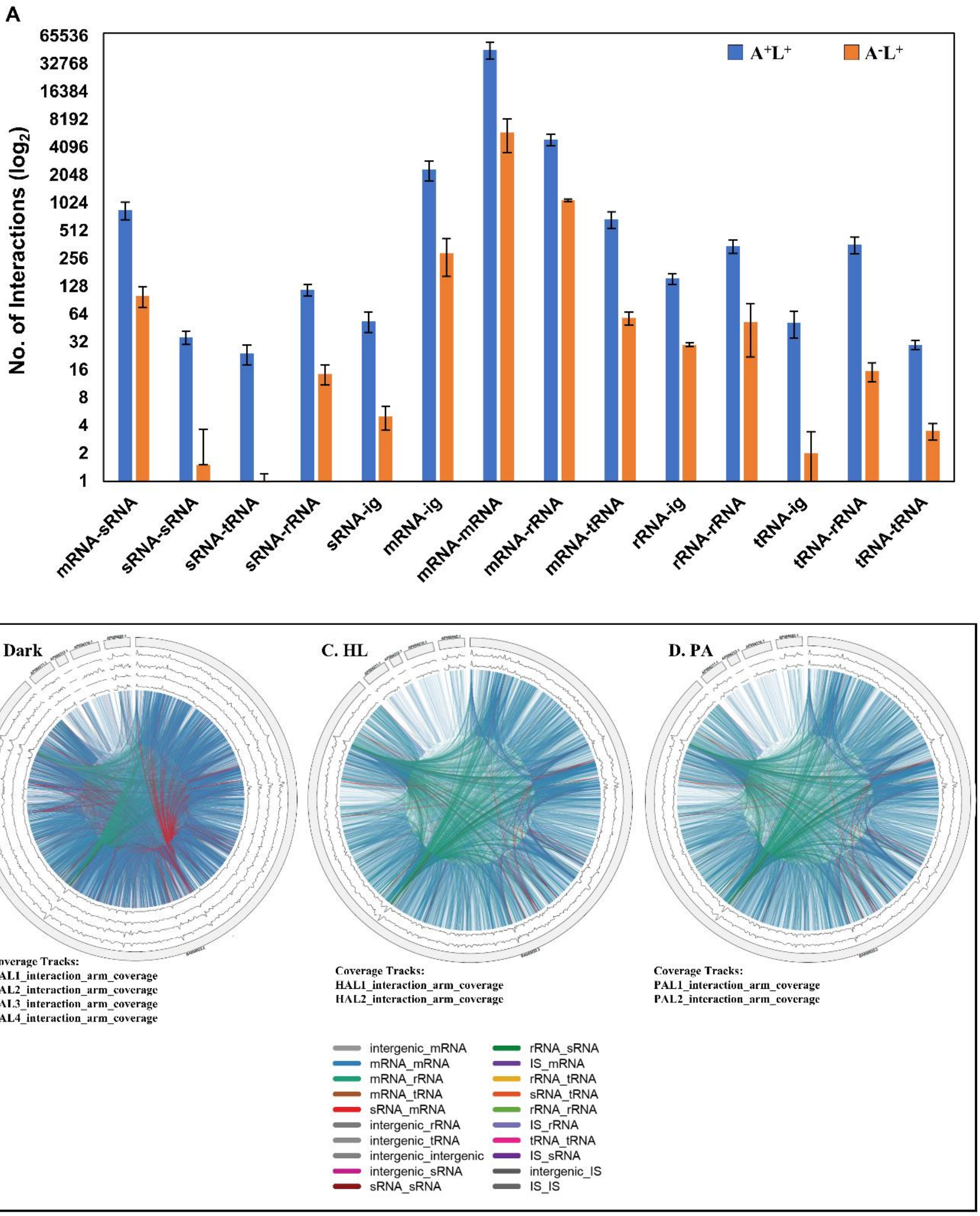


**Figure S2: Global RNA–RNA interaction landscapes in *Synechocystis* sp. PCC 6803.**

**A:** RNA–RNA interactions were filtered independently in each biological replicate using an RNANue adjusted p-value (padj_value) < 0.1. Bars represent the mean number of significant interactions across the AMT-treated and ligated ($A^{+}L^{+}$) and uncrosslinked and ligated ($A^{+}L^{-}$) biological replicates for each interaction category. Error bars indicate standard deviation.
**B–D:** Global circos plots interaction maps containing interactions between all RNA transcript types represented in the interaction dataset under different conditions. The outermost ring denotes the chromosome and four plasmids, with interaction-arm coverage represented by the inner genomic tracks derived from each biological replicate. The arcs indicate the interacting RNAs at their corresponding positions and color-coded for different interaction groups. This figure extends Figure 4.

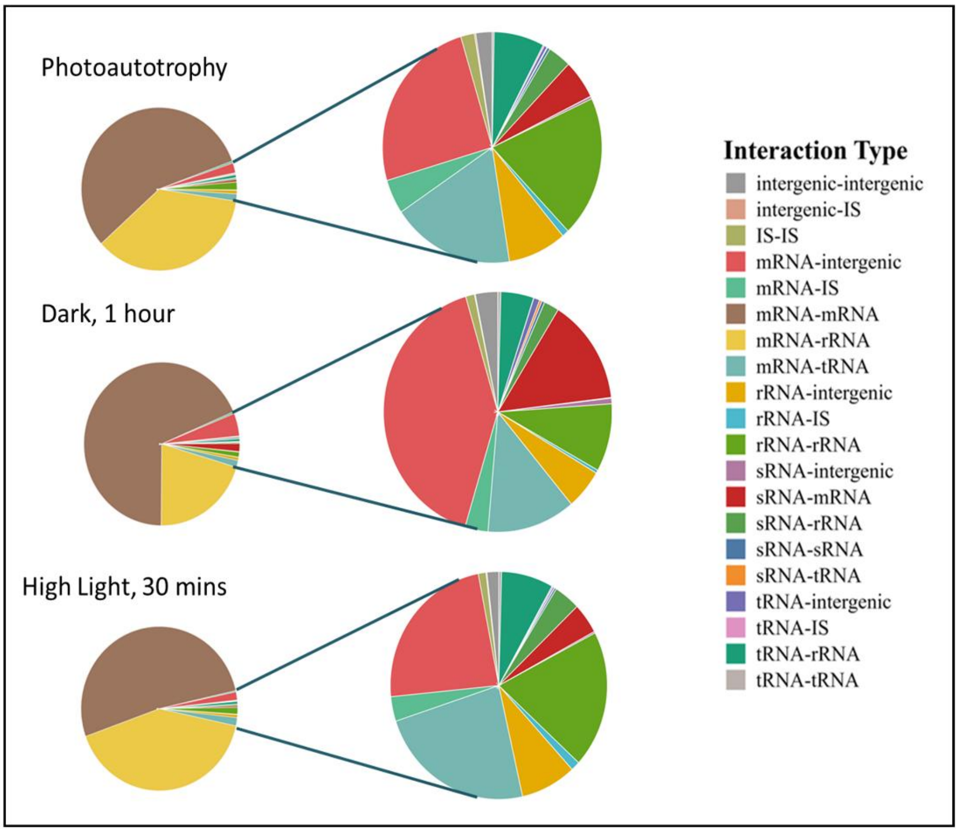


**Figure S3: LIGR-NUE analysis of *Synechocystis* RNA-RNA interactome**

(A) The pie-of-pie graphs represent counts of interactions among different transcript types across the given conditions. The main pie illustrates the number of interactions for all interaction types, while the inset pie provides an expanded view of all other interactions except mRNA-rRNA and mRNA-mRNA. This figure extends Figure 4.

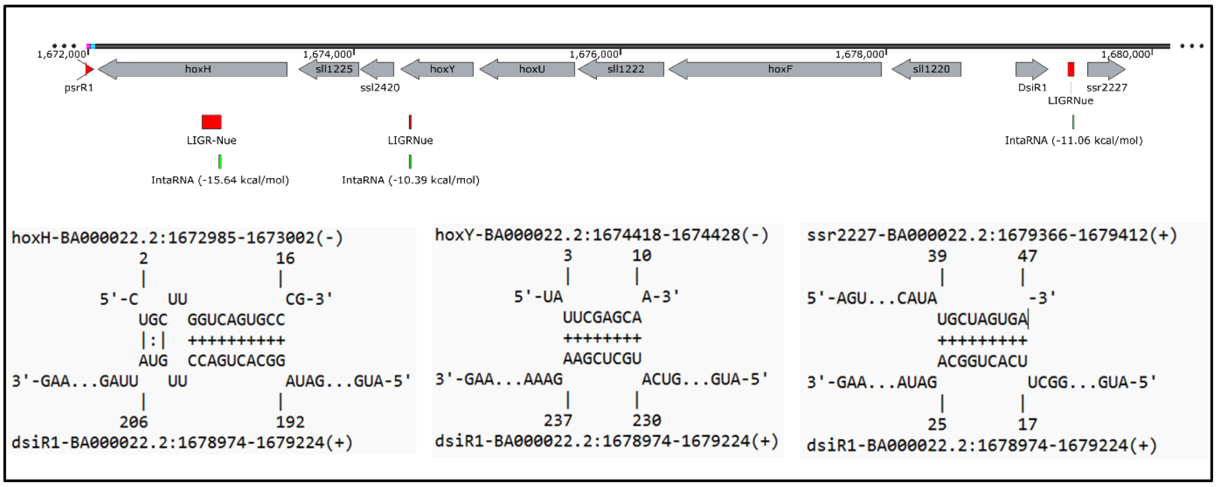


**Figure S4: Snapshot of DsiR1 interactions with adjacent genes *hoxH*, *hoxY* and *ssr2227*: Top:** Genomic organization of the *hox* locus (encodes bidirectional hydrogenase), *DsiR1* and *ssr2227* (encodes putative transposase). Red bars indicate interaction fragments detected by LIGR-Nue, whereas green bars denote interaction regions predicted by IntaRNA; predicted interaction energies are shown in brackets. **Bottom:** IntaRNA-predicted base-pairing structures between *DsiR1* and the respective target regions of *hoxH*, *hoxY*, and *ssr2227*. The predicted interaction energies are −15.64, −10.39, and −11.06 kcal/mol, respectively.

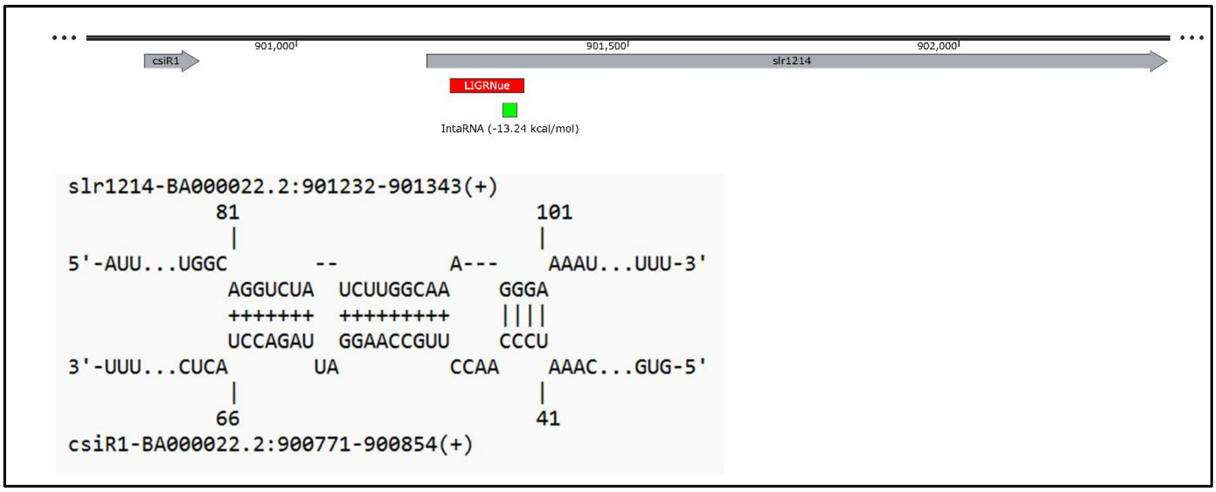


**Figure S5.** Snapshot representation of the predicted base-pairing interaction between the sRNA CsiR1 and slr1214, which encodes a response regulator. **Top:** Genomic organization showing the location of *csiR1* upstream of *slr1214*. The red bar indicates the interaction fragment detected by LIGR-Nue, while the green bar represents the interaction region predicted by IntaRNA. **Bottom:** IntaRNA-predicted base-pairing structure between CsiR1 and *slr1214*, with interaction energy of −13.24 kcal/mol.

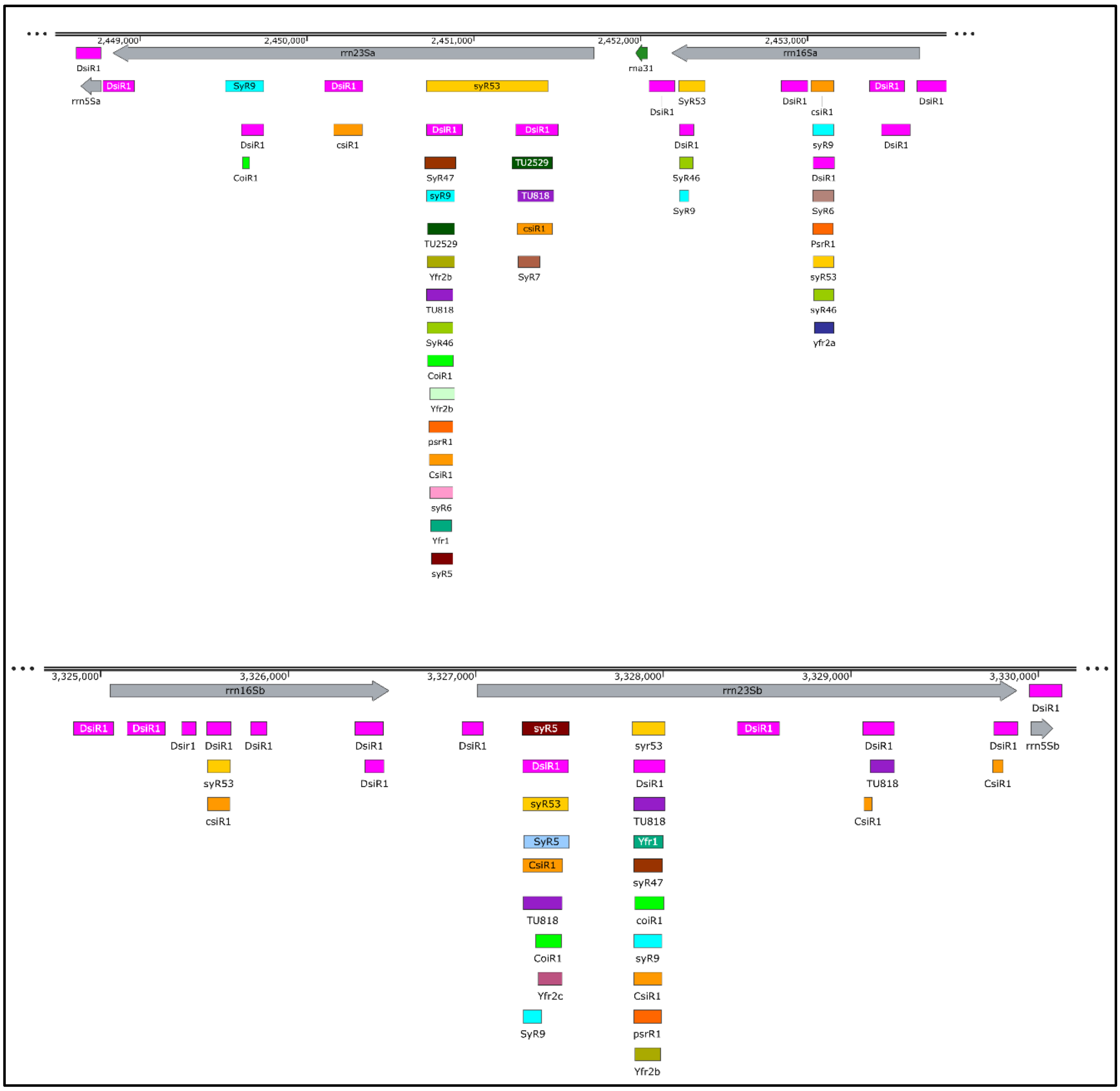

**Figure S6:** Snapshot showing interactions of different sRNAs with the rRNA genes rrn23Sa, rrn16Sa, rrn23Sb, rrn5Sa, rrn5Sb and rrn16Sb at specific positions. All sRNAs are observed to interact at particular conserved locations within these rRNA regions. The figure also highlights multiple interaction sites of DsiR1with rRNAs exhibiting interactions at several locations under dark conditions.
